# Sodium butyrate is incorporated into central metabolism in fly head while inducing oxygen consumption increase

**DOI:** 10.1101/2024.07.18.604059

**Authors:** Annika Müller-Eigner, Benedikt Gille, Frederik Dethloff, Chen Meng, Christina Ludwig, John T Heiker, Patrick Giavalisco, Shahaf Peleg

## Abstract

Butyrate has been proposed as a drug therapy by acting as a KDAC inhibitor and elevating protein acetylation, in particular on histones. Nonetheless, recent studies suggest that tissues such as the gut can utilize butyrate as a metabolite. We have previously shown that the addition of butyrate induces a rapid increase of oxygen consumption in whole *Drosophila melanogaster* heads. Here we show that while head oxygen consumption is increased by the addition of butyrate, no apparent changes are observed on the proteome and acetylome. Instead, we show that butyrate is metabolized and incorporated into the tricarboxylic acid cycle (TCA) cycle. Collectivity our data supports the notion that the therapeutic benefits of acute butyrate treatment may be also mediated by improving metabolic rates, rather than solely targeting the epigenome or acetylome.

## Introduction

Chromatin remodeling is a key mechanism, which enables living cells to react and adjust to changes in the environment. One core mechanism to modulate the chromatin is the modification of histones, a set of proteins that tightly interacts and packages the DNA (Kornberg, 1974; Luger et al., 1997; Kornberg and Lorch, 1999). Modification of histones by small metabolites lead to alterations in the binding between histone and DNA, resulting in fine-tuning of transcription (Strahl and Allis, 2000; Wang et al., 2022). One of the most studied modifications of histone proteins is histone acetylation, which is commonly associated with reduced histone-DNA interaction that leads to increased transcription activity (Berger, 1999; Rice and Allis, 2001; Cedar and Bergman, 2009; Choudhary et al., 2014).

Dysregulation of chromatin remodeling during disease and aging has attracted considerable attention in the recent two decades. For example, the observation that histone acetylation levels are altered during aging and maladies such as neurodegeneration and Alzheimer’s disease has led to identifying histone acetylation as an important target (Peleg et al., 2010). Indeed, the inhibition of histone deacetylases (HDACs) by drugs such as suberoylanilide hydroxamic acid (SAHA), butyrate and others has been shown to improve age-associated memory formation deregulation, neurodegeneration, muscle function and related metabolism, heart diastolic dysfunction and others in mice (Paskova et al., 2013; Park and Sohrabji, 2016; Fernando et al., 2020). Therefore, a strong link between inhibiting HDAC, increased histone acetylation and disease therapy was suggested.

Nonetheless, there is growing evidence in recent years that many non-histone proteins are acetylated as well (Friedmann and Marmorstein, 2013; Choudhary et al., 2014; Schölz et al., 2015; Drazic et al., 2016; Imhof and Peleg, 2016). These proteins are involved in various cellular pathways. Of note, HDACs were shown to be able to deacetylate many of these non-histone proteins and thus were more accurately termed Lysine Deacetylases (KDAC) (Imhof and Peleg, 2016). As such, it is possible that KDAC inhibitors (KDACi) could impact a myriad of non-epigenetic pathways.

Previously, we have shown that sodium butyrate (SB) induces rapid and transient oxygen consumption increase in the heads of *Drosophila melanogaster* analyzer (Peleg et al., 2016; Becker et al., 2018; Dietz et al., 2019) while other work showed similar induction in colon-derived cell lines (Bekebrede et al., 2021). The rapid increase was detected already after several minutes, suggesting that it may be too rapid to be dependent on new gene transcription following protein translation that is mediated by histone acetylation changes (Schwanhäusser et al., 2011). Interestingly, previous studies have shown that butyrate can induce increased non-histone protein acetylation (Schölz et al., 2015). As many of these acetylated proteins are known to be metabolic enzymes, we hypothesized that butyrate may cause an increase in the acetylation levels of metabolic proteins, which may underlie the observed rapid increase in oxygen consumption rate (OCR). We expected that butyrate will impact the levels of metabolites associated with mitochondrial oxygen consumption.

## Results

### Oxygen consumption assessment in fly heads

We have previously measured the impact of SB on OCR in a Seahorse X24 analyzer (Peleg et al., 2016; Becker et al., 2018; Dietz et al., 2019). In the present study, we first re-assessed the kinetics of SB-induced OCR increased using the new Seahorse XFe24 analyzer with improved sensor capacity. Furthermore, we improved the pH adjustment of media containing SB/vehicle (see methods). We also tested lower and higher concentrations of SB (3mM, 30mM). Using the improved protocol, we observed that compared to previously published data, 15 mM SB induced an even more rapid OCR increase, peaking as early as 22.5 minutes (4^th^ measurement compared with 6^th^ measurement in (see Becker et al., 2018)) (Figure 1 and S1B). Additionally, 30mM SB induced a stronger and prolonged OCR increase (Figure 1 and S1C), whereas, 3mM SB treatment was insufficient and did not result in an OCR increase (Figure 1 and S1A). Similarly, under the improved protocol, vehicle group showed relatively more stable OCR levels throughout the experiment (Figure 1), compared to our previous work where the vehicle measurements displayed a mild and transient increase in OCR. Collectively, the improved protocol supported the notion of earlier impact of SB on OCR alongside an improved control group.

**Figure 1:**
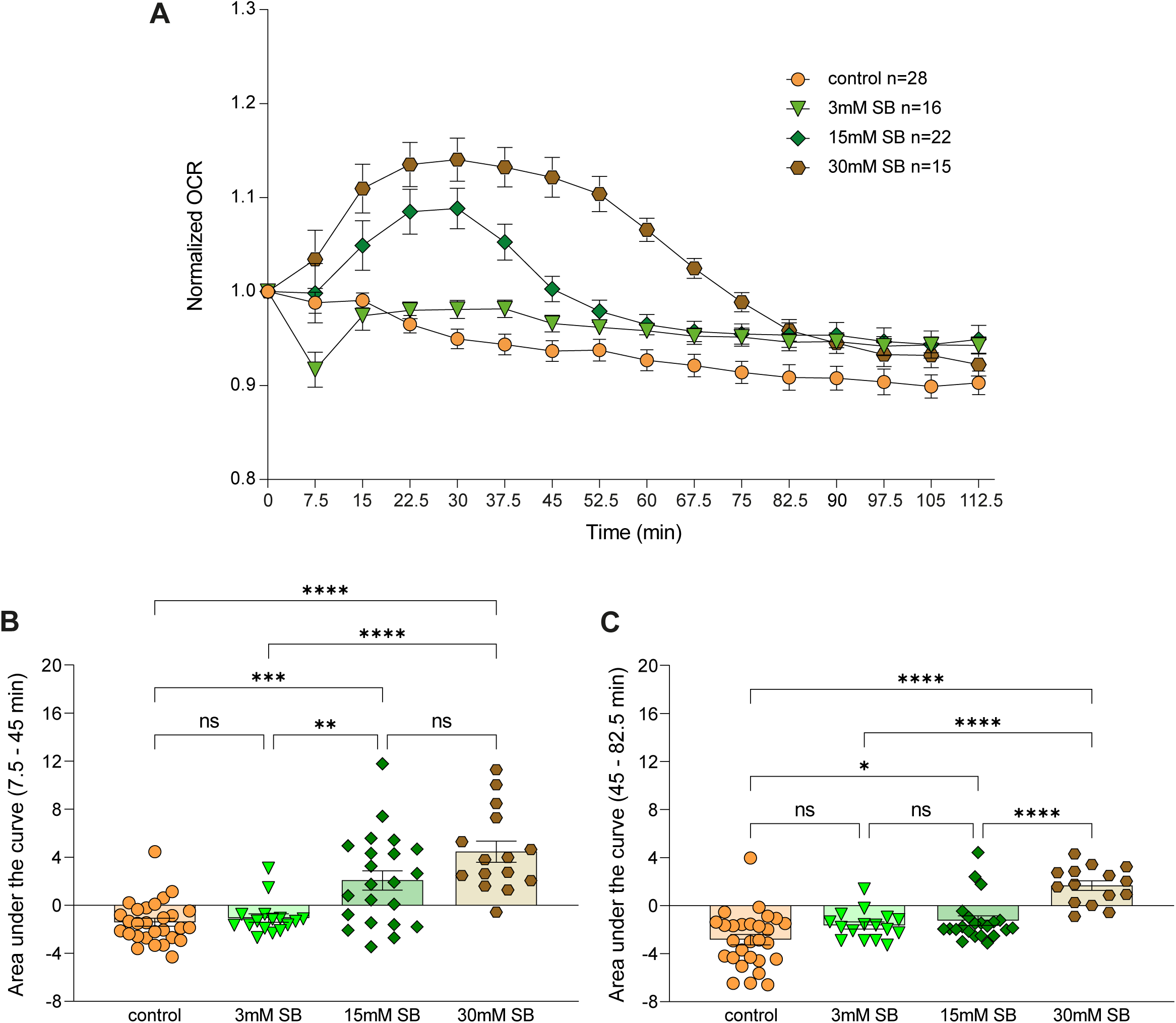
Sodium butyrate rapidly increase oxygen consumption in a dosage dependent manner. (A) Quantification of oxygen consumption levels upon addition of SB. Time zero refers to the last measure prior to the addition of vehicle/SB. (B+C) Area under the curve for each condition plotted in a scatter plot for the 2-7 (7.5 – 45 minutes) measurements (B) after the injection and 7-12 (45 – 82.5 minutes) measurements (C) after the injection. Control (orange, n=28), 3 mM SB (light green, n=16), 15 mM SB (dark green, n=22), 30 mM SB (brown, n= 15). Error bars indicate standard error of mean in all graphs. One-way ANOVA, Tukeýs post-hoc test, ns p>0.05; * p≤0.05; ** p≤0.01; *** p≤0.001; **** p≤0.0001. Detailed statistical information is presented in Supplementary Table 1 and 2.

We next assessed the impact of SB treatment on the fly head proteome and acetylome while focusing on potential alterations during 20 minutes of 15mM SB treatment.

It was hypothesized, that sodium butyrate, as a KDACi, alters non-histone protein acetylation in fly heads, which leads to increased oxygen consumption, and that the observed phenotype is not caused by any transcriptional changes due to the short treatment window of only 20 minutes (Schwanhäusser et al., 2011). As expected, our proteome comparison between vehicle and 15mM SB of 2827 detectable proteins shows no significant abundance change in flies’ heads after 20 minutes of incubation with SB or vehicle solution (Figure 2A). Specifically, we detected no significant changes in the respiratory complexes 1-4 (Figure S2) and key metabolic enzymes in the TCA cycle metabolism (Figure S3).

**Figure 2:**
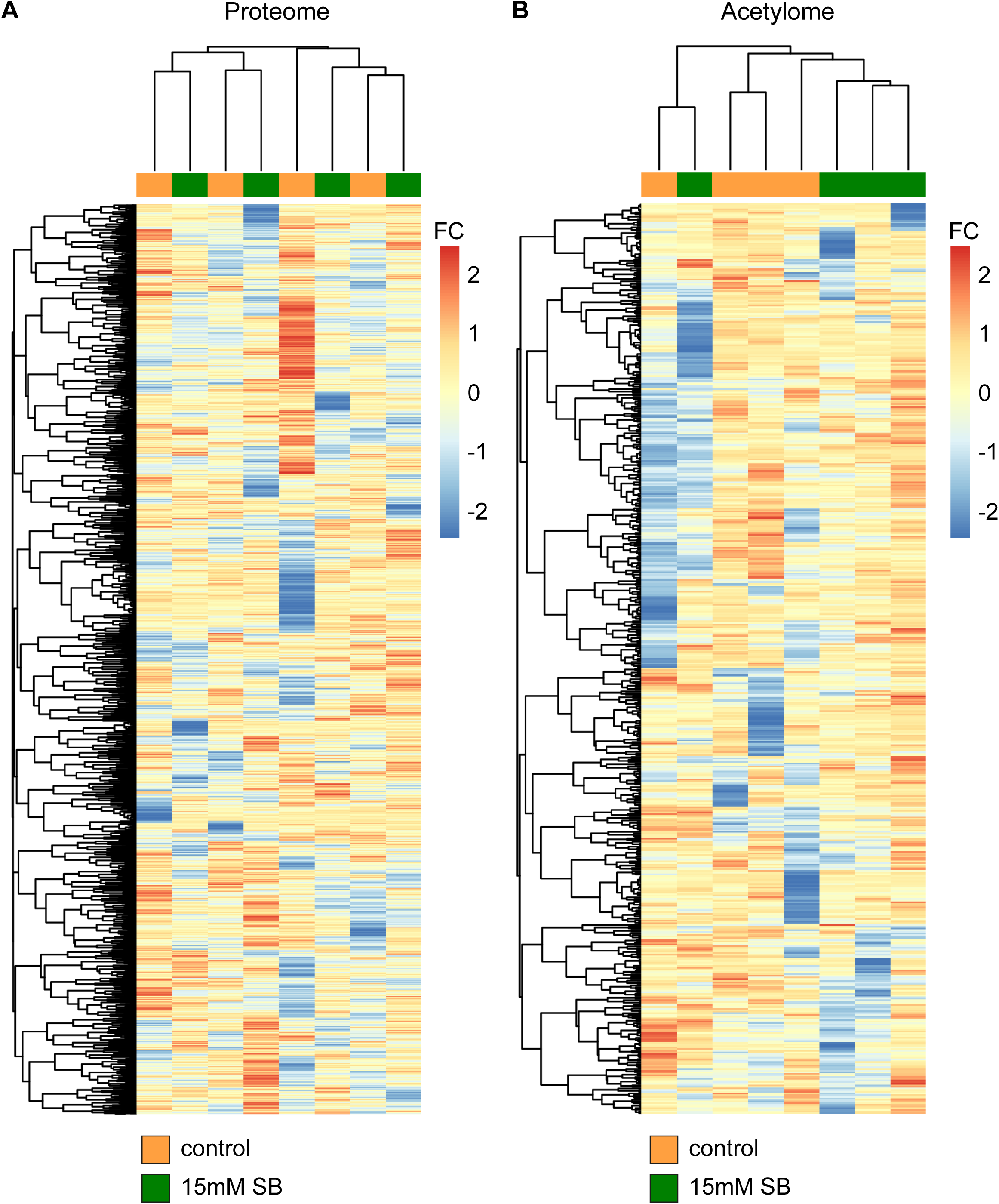
Sodium butyrate does not impact the levels of proteins and acetylation of proteins after 20 minutes. (A) Heatmap showing comparison of 2827 detected proteins between control (orange, n=4) and 15 mM SB (green, n=4) treated fly heads after 20 minutes. (B) Heatmap comparing 557 detected acetylation sites of 267 proteins between control (orange, n=4) and 15 mM SB (green, n=4) treated fly heads after 20 minutes. FC (fold change).

Nonetheless, SB has been previously shown to be a KDAC inhibitor that affects protein acetylation, which is more rapid and dynamic compared with overall protein abundance alterations (Donohoe et al., 2011; Mollica et al., 2017; Li et al., 2018; Li et al., 2019) . Dynamic protein acetylation itself has been shown to impact metabolic regulation (Choudhary et al., 2009; Zhao et al., 2010; Shang et al., 2022). Therefore, we next measured the protein acetylome after SB/vehicle treatment at 20 minutes. Interestingly, no significant alterations were found among 267 different proteins harboring 557 acetylation sites (Figure 2B). Collectively, we found no impact of SB neither on protein levels nor on protein acetylation at the peak of OCR increase in flies’ heads.

As butyrate does not impact protein abundance and protein acetylation, at least at the 20 minutes mark, we wondered if the heads can utilize butyrate as a metabolite. Previous work showed that SB is used as a metabolite in colonocytes and influences mitochondrial function, e.g., in the liver (Donohoe et al., 2011; Mollica et al., 2017). As such, we hypothesized that butyrate could be utilized as an energy source in fly heads.

Firstly, we tested metabolites that are known to affect the electron transport chain and hence the OCR. In a similar experimental design described for the SB treatment, we added pyruvate and proline as positive controls to fly heads. Indeed, treatment of fly heads with pyruvate and proline, two metabolites that are known substrates for fly mitochondria, revealed kinetics similar to the responses observed after SB treatment (Figure 3). The similarity of the kinetic of pyruvate/proline and SB suggests that SB might be used as mitochondrial substrate in the heads.

**Figure 3:**
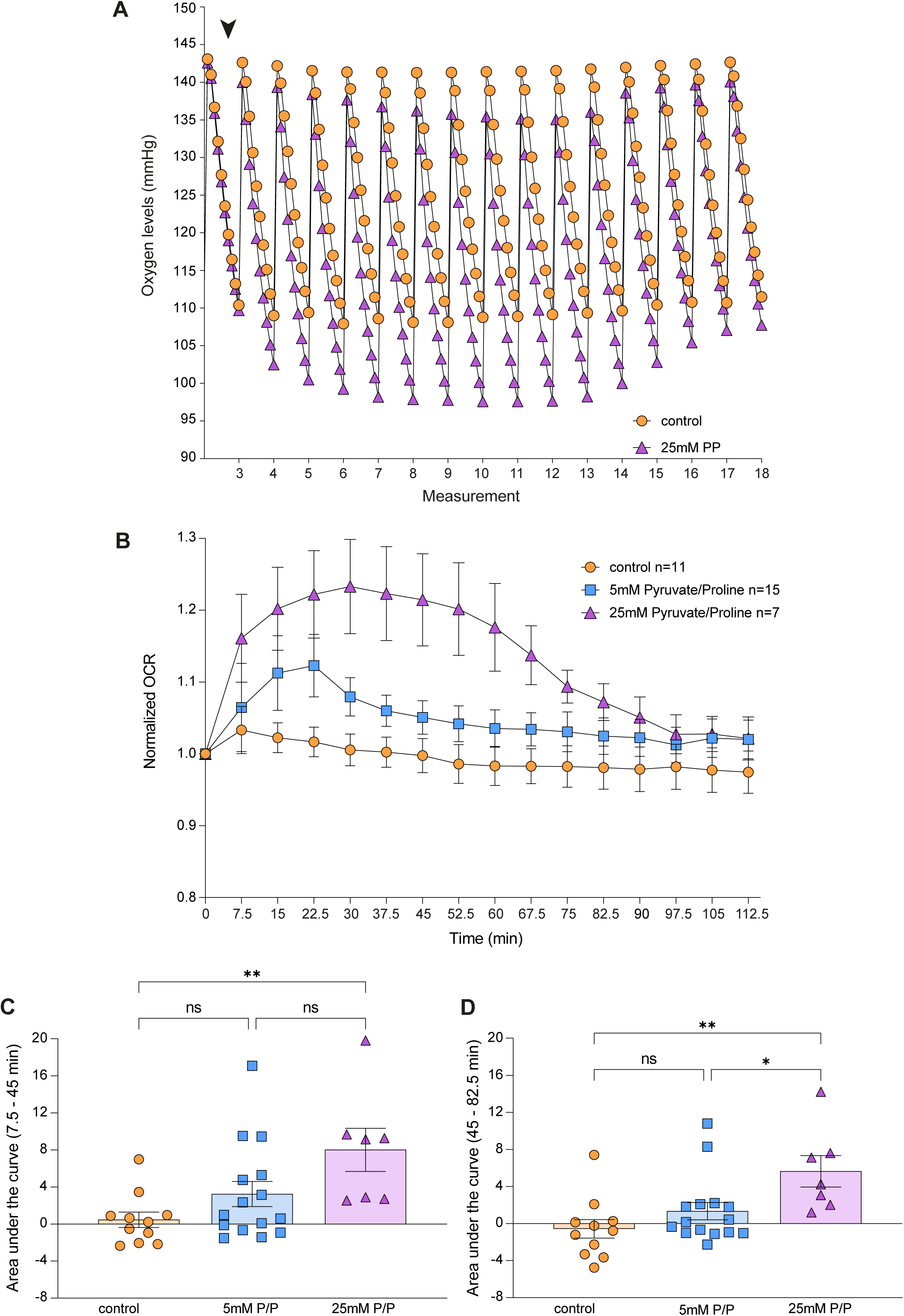
Pyruvate and proline rapidly and transiently increase oxygen consumption in fly heads. (A) Representative tracing for oxygen level changes in control and 25 mM pyruvate and proline (PP)-treated fly heads. Each measurement lasted 2 minutes and consisted of 10 sub-measurements (ticks). The last measurement before (3) and the measurements after injection (4-18) are shown. The arrowhead indicates the addition of PP or control. (B) Quantification of OCR based on (A). (C+D) Area under the curve for each condition plotted in a scatter plot for the (C) 2-7 (7.5 – 45 minutes) measurements or (D) 7-12 (45 – 82.5 minutes) measurements after the injection. Control (orange, n=11), 5 mM PP (blue, n=15), 25 mM PP (purple, n=7). One-way ANOVA, Tukeýs post hoc test, ns p>0.05; * p≤0.05; ** p≤0.01. Error bars indicate ±S.E.M in all graphs. Detailed statistical information is presented in Supplementary Table 4 and 5.

To test this hypothesis, we incubated fly heads in a buffer containing ^13^C_4_-labeled 15 mM SB and the assessed incorporation of labeled carbon atoms into metabolites by using liquic chromatography-mass spectrometry (LC-MS) at three distinct time points (0 min, 20 min, and 60 min). Of note, fly heads were continuously exposed to ^13^C_4_-labeled 15 mM SB for the entire duration of each experiment. As expected we found substantial increase in the pool sizes of butyric acid, butyryl coenzyme A (CoA), and 3-hydroxybutyryl-CoA (Figure 4) showing nearly 100 % incorporation of heavy labeled carbons (Figure S4 and Figure S5A-C). Additionally, we observed increased ^13^C_2_ labeling of acetyl-CoA over time (Figure S6 and Figure S5D). In line with these results, we also observed a substantial increase in stable isotope labeling in all detected TCA metabolites (Figure 5 and Figure S5E-M), indicating that the carbons from the butyrate have been rapidly incorporated into the TCA cycle specifically via β-oxidation to acetyl-CoA. Notably, the overall incorporation of ^13^C_4_ continued to increase from 20 minutes to 60 minutes (Figure 5). Lastly, we did not observe ^13^C incorporation in this time frame into other metabolites such as the amino acids arginine, aspartate, glutamate and proline (Figure S7).

**Figure 4:**
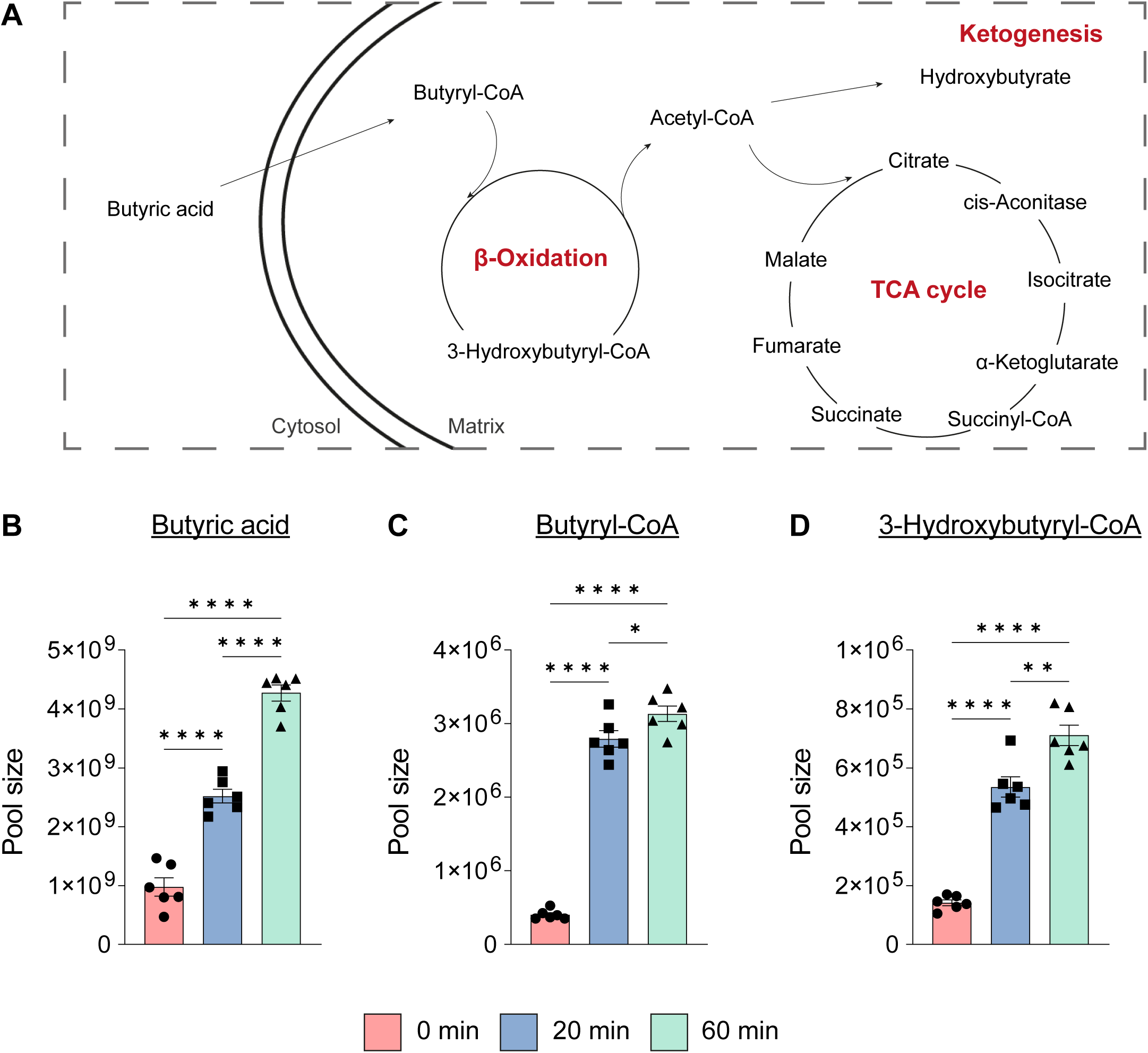
Sodium butyrate is rapidly incorporated into butyric derivatives. (A) Schematic overview of key metabolic pathways (red) involving the measured metabolites. (B-D) Changes in pool size in response to ^13^C_4_-labeled SB treatment in fly heads over time (red, 0 min; blue, 20 min; green, 60 min). n=6 for all metabolites. Error bars indicate ±S.E.M in all graphs. One-way ANOVA, Tukeýs post-hoc test, ns p>0.05; * p≤0.05; ** p ≤ 0.01; *** p≤0.001; **** p≤0.0001. Detailed statistical information is presented in Supplementary Table 6.

**Figure 5:**
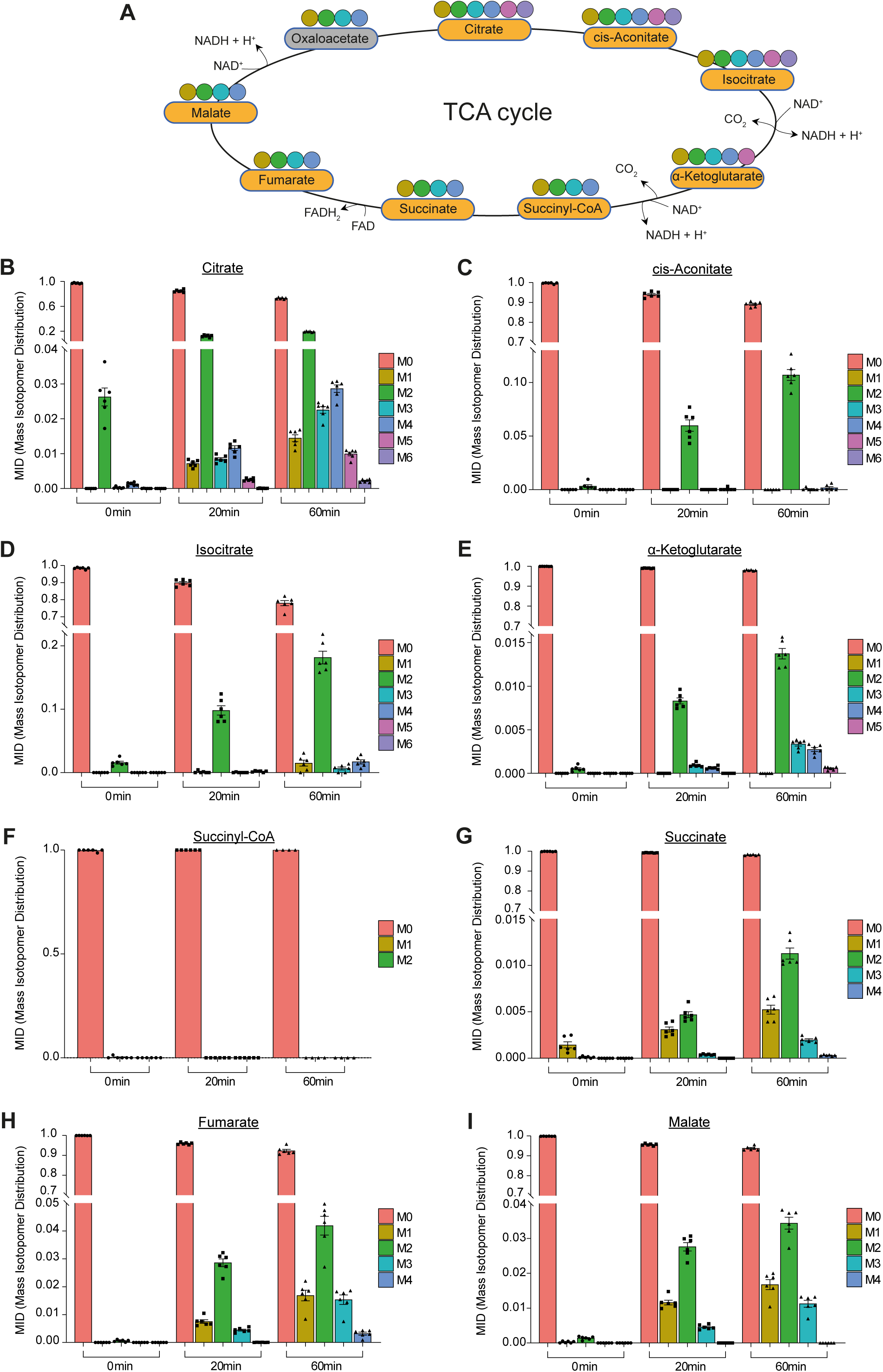
Butyrate is rapidly incorporated into TCA cycle metabolites. (A) Schematic overview of the TCA cycle with measured metabolites (orange) and their specific carbon atoms (colored circles). (B-I) Analysis of Mass Isotopomer Distribution (MID) in TCA cycle metabolites after ^13^C_4_-SB treatment to determine its incorporation over time by mass spectrometry. n=6 for all metabolites and time points, except for succinyl-CoA 60 min n=4. M0 unlabeled mass of isotope, M+n native metabolite mass + number of isotopically labeled carbons. Error bars indicate ±S.E.M in all graphs. Detailed statistical analysis was performed using one-way ANOVA, Tukeýs post-hoc test, results are presented in Supplementary Table 7.

## Discussion

In this work, we tested the impact of SB on molecular metabolism processes that might explain the rapid increase in whole fly head OCR. Firstly, we used an improved method to measure OCR in response to SB treatment. The new Seahorse analyzer, manual OCR calculation (see methods) and improved pH control have led to a more stable result for vehicle group measurements, compared with previously published data (Becker et al., 2018). Furthermore, the improved method revealed an earlier increase of OCR in response to SB feeding, namely already after 3-4 measurements (22,5-30 minutes), which supported the notion that the OCR increase is independent of histone-mediated transcription and following translation (Schölz et al., 2015; Peleg et al., 2016). Additionally, we showed that the increased OCR is SB is dose-dependent. For example, using higher 30mM SB concentration results in stronger and prolonged OCR increase in fly heads. We suggest that molecular studies could benefit from using increased dosage of SB that may assist uncovering the impacted molecular pathways, as the magnitude of response is elevated.

Notably, we observed that similarly to SB, pyruvate and proline induce similar increased OCR kinetics. Interestingly, both responses are transient and are back to normal OCR levels after over one hour. The reason for this is not clear and should be further investigated. It is evident that, at least with SB, sufficient SB is still present in the buffer after 60 minutes (Figure S8). However, it is possible that feedback loops aimed to control mitochondrial respiration are active to counter SB’s impact on the OCR. More work is needed to understand the mechanism that reset the OCR levels back to their basal state.

In line with our initial hypothesis, we found no changes in protein abundance after 20 minutes of SB treatment. This is supported by the fact that such time frame is likely too short for the transcription and translation of proteins to increase the OCR (Schwanhäusser et al., 2011). However surprisingly, in contrast to our initial hypothesis, we also observed no significant alterations in protein acetylation in this time frame. Protein acetylation was shown to modulate metabolic activity, and acetylation itself is rapid and dynamic (Zhao et al., 2010; Schölz et al., 2015; Shang et al., 2022). While it is possible that a broader mass spectrometric coverage would reveal protein acetylation changes, our data support the notion that SB, at least within the measured time frame, does not induce increased acetylation that may be linked with the increased OCR. This is important as previous studies tested short-term impact of SB in the brain, for example on memory formation enhancement, and linked it with acetylation increase (Fernando et al., 2020). However, our data suggest that SB may in fact act as a metabolic boost and that such a boost may be the underlying benefit for previously described therapeutic attributes of SB.

Our results further support the notion that butyrate is rapidly incorporated specifically into β-oxidation and TCA cycle metabolites. We observed a clear ^13^C incorporation in the TCA cycle metabolites. Notably, the measurement intensity for succinyl-CoA was low, while oxaloacetate was not detected at all, which might indicate lower stability of these metabolites, therefore making it more difficult to detect ^13^C. Nonetheless, our data clearly shows a specific and gradual accumulation of ^13^C in acetyl-CoA and TCA metabolites. In fact, already at so-called time point 0, we observed ^13^C incorporation. This might be attributed to the fact that even the short exposure to ^13^C_4_ labeled butyrate, while collecting the samples, is sufficient to cause ^13^C incorporation into the TCA metabolites. Overall, this data shows a correlation between fueling the oxidation and TCA cycle in a similar time frame to increased OCR in fly heads.

In summary, we show that SB rapidly incorporates to central metabolic pathways in fly heads while, at least within the studied time frame, causing no visible impact on protein acetylation. Our data supports the notion that the role of SB as a therapy mainly via inhibiting KDAC to cause increased protein acetylation should be revisited. Instead, we rationalize that SB may act, at least acutely, as a transient metabolic boost and thus lead to improved memory performance as previously described. Nonetheless, more causal evidence is needed to tightly link the observed OCR increase induction by SB and rapid metabolic alterations in the context of whole tissue.

## Methods

### Fly Strains

The *Drosophila melanogaster* strain *w^1118^* (Canton) was maintained and cultivated at 25 °C with 60% humidity and a 12 h light/ 12 h dark cycle on standard fly food (In 11L dest. H2O, 97,5g Agar, 975g cornmeal (biogewinner, order nr. 544) 112,5g soy flour GMO free (Bakeryshop), 225g dry yeast (Mühle Schlingemann, order nr.: 110115), 975g melasse (Grafschafter Krautfabrik, order nr.: 1939), 487,5g malt (Diamalt, CSM Deutschland GmbH), 31,125g Nipagin (Roth Order Nr. 3646.4) and acid mix ((168,37ml H2O + 6,64ml Phosphoric acid (Sigma, Order nr.:345245), 46,24mL Propionic acid (Sigma, Order nr.: 81910)) and transferred to new vials with fresh fly food at different time periods. In order to select flies according to gender, flies were briefly sedated with CO_2_. Male flies were selected for the study at the age of 7-9 days.

### Oxygen consumption measurements

Oxygen consumption measurements were done using Seahorse XFe24 Analyzer and the corresponding XFe24 island plates as previously described (Peleg et al., 2016) with the following modifications:

Solutions were freshly prepared on experimental day and pH was adjusted to 7.15 (buffer) and 6.95 (SB) with NaOH or HCl at experimental temperature of 31 °C. The plate wells and the injection ports were inspected for air bubbles, which were removed using a pipette if necessary.

The automatic FIXED algorithm, which was used for the XF24 machine (Dietz et al., 2019), is absent in the new XFe24 machine and therefore OCR was analyzed by calculating the decrease in oxygen level by dividing tick 5 by tick 2 and normalized to the last basal measurement before injection (third total measurement).

### Plate preparation for omics

Plate preparation for proteome/acetylome analyses was as done as described preparation for the Seahorse XFe24 Analyzer with the following additions (Dietz et al., 2019). The plate was incubated at 31 °C for 20 minutes to reflect the three basal measurements during the OCR experiments. Afterwards, vehicle or SB solution was carefully added to fly heads and an incubation step at 31 °C of 20 minutes or 60 minutes followed. Subsequently, plates were removed from the incubator and supernatant was discarded. The wells were washed with water and immediately frozen by using liquid nitrogen. Fly heads were removed on dry ice from plates by using forceps and stored in reaction tubes at – 80 °C until further analysis. 16 fly heads corresponded to one biological sample (n) for metabolome analysis, whereas 1000 fly heads were needed for proteome and acetylome analysis.

### Proteome and acetylome analysis

High-resolution mass spectrometry was used to identify potential lysine acetylation sites on proteins and acetylation changes in fly heads as previously described (Peleg et al., 2016) .

Briefly, 100 mg of frozen fly heads were homogenized in 300 µl lysis buffer as previously described (Peleg et al., 2016) . To reduce disulfide bonds, samples were treated with the reducing agent dithiothreitol (DTT), followed by a treatment with iodoacetamide (IAA) to alkylate free cysteines.

Samples were digested for 5 h with Lys-C and with trypsin. The eluted peptide concentration was measured by NanoDrop One^C^ spectrophotometer and equal concentration of peptides were incubated with 45 µl anti-acetyllysine antibody (ICP0388, Immunechem) overnight at 4 °C. Beads were washed four times with phosphate-buffered saline (PBS)/Tween 0.1% and then four times with PBS. Acetylated peptides were eluted from beads with 125 µl 0.1% Trifluoroacetic acid (TFA). For proteome analysis 20 µl of each sample were taken before enrichment for acetylated lysines and were diluted with 80 µl 0.1% TFA.

LC-MS/MS measurements were performed as described previously (Müller-Eigner et al., 2022). Briefly, for full proteome measurement, 0.25 µg purified peptides were injected into an Ultimate 3000 RSLCnano system coupled to a Q-Exactive HF-X mass spectrometer (Thermo Fisher Scientific), while for acetylome measurements the complete peptide sample derived after acetylome enrichment (concentration unknown) was injected. The Q-Exactive HF-X mass spectrometer was operated in the data-dependent acquisition and positive ionization mode using a 120 minute linear LC gradient. Peptide identification and quantification were performed using MaxQuant (version 1.6.3.4). MS2 spectra were searched against the Uniprot *Drosophila melanogaster* proteome database (UP000000803, 21923 entries, download at 22.3.2019). For full proteome analyses, carbamidomethylated cysteine was set as fixed modification and oxidation of methionine and N-terminal protein acetylation as variable modifications. For the acetylome data, additionally Lysine acetylation was set as variable modifications.

Protein abundances were calculated using the label-free quantification algorithm from MaxQuant (Cox et al., 2014). The intensities of lysine-acetylated peptides from the “modificationSpecificPeptides.txt” table were used to quantify the relative abundances of acetylation site. Student’s t-test was used to identify the differentially expressed proteins and lysine-acetylated peptides between control and SB. The resulting p values were adjusted by the Benjamini–Hochberg algorithm the FDR (Benjamini and Hochberg, 1995). For heatmap generation, the intensity matrix was scaled row-wise. Correlation distance and Euclidean distance were used to generate dendrograms on rows and columns respectively. The statistical analyses were performed using R 4.1.0 (R Core Team, 2021).

### Metabolome analysis

Heavy-labeled sodium butyrate-^13^C_4_ was used for all metabolome-experiments. Semi-targeted liquid chromatography-high-resolution mass spectrometry (LC-HRMS) analysis of short chain fatty acyl coenzyme A species (Acyl-CoAs) was applied.

The LC-HRMS analysis of Acyl-CoAs was performed using an adapted protocol (Abrankó et al., 2018).

In brief, the dried metabolite extract was re-suspended in 50 µl of LC-MS-grade water (Optima-Grade, Thermo Fisher Scientific). After 15 minutes incubation on a thermomixer at 4 °C and a 5 minutes centrifugation at 16,000 x g at 4 °C, the cleared supernatants were transferred to glass autosampler vials with 300 µl glass inserts (Chromatography Accessories Trott, Germany).

For the LC-HRMS analysis, 1 µl of the sample was injected onto a 30 x 2.1 mm BEH Amide UPLC column (Waters) with 1.7 µm particle size. The gradient elution ran in a quaternary Vanquish Flex LC system (Thermo Fisher Scientific) with a flow rate of 500 µl/min using a quaternary buffer system consisting of buffer A 5 mM ammonium acetate (Sigma) in LC-MS-grade water (Optima-Grade, Thermo Fisher Scientific); buffer B consisted of 5 mM ammonium acetate (Sigma) in 95% acetonitrile (Optima-grade, Thermo Fisher Scientific); buffer C consisted of 0.1% phosphoric acid (85%, VWR) in 60% acetonitrile (acidic wash) and buffer D of 50% acetonitrile (neutral wash). The column temperature was set to 30 °C, while the LC gradient was: 85% B for 1 minutes, 85-70% B 1-3 minutes; 70-50% B 3 – 3.2 minutes; holding 50% B till 5 minutes; 100% C 5.1 – 8 minutes, 100% D 8.1 - 10 minutes; followed by re-equilibration with 15% A and 85% B 10.1 – 13 minutes. The mass spectrometer (Q-Exactive HF, Thermo Fisher Scientific) was operating in positive ionization mode recording the mass range m/z 760-1800. The heated electrospray ionization (ESI) source settings of the mass spectrometer were set to: Spray voltage 3.5 kV, capillary temperature 300 °C, sheath gas flow 50 AU, aux gas flow 15 AU at a temperature of 350 °C and the sweep gas to 3 AU. The RF-lens was set to a value of 55.

The identity of acetyl-CoA and malonyl-CoA was validated by authentic ^13^C labeled reference compounds, which were run before. Other Acyl-CoAs annotations were validated by using *E. coli* reference material matching exact mass and reporter ions from PRM experiments. Peak areas of [M + H]^+^ ions and corresponding isotopomeres were extracted using a mass accuracy (<5 ppm) and a retention time tolerance of <0.05 minutes.

### Semi-targeted LC-HRMS analysis of amine-containing metabolites

The LC-HRMS analysis of amine-containing compounds was performed using an adapted benzoylchlorid-based derivatization method (Wong et al., 2016).

In brief: The polar fraction of the metabolite extract was re-suspended in 200 µl of LC-MS-grade water (Optima-Grade, Thermo Fisher Scientific) and incubated at 4 °C for 15 minutes on a thermomixer. The re-suspended extract was centrifuged for 5 minutes at 16,000 x g at 4 °C and 50 µl of the cleared supernatant was mixed with 25 µl of 100 mM sodium carbonate (Sigma), followed by the addition of 25 µl 2% [v/v] benzoylchloride (Sigma) in acetonitrile (Optima-Grade, Thermo Fisher Scientific). Samples were vortexed and kept at 20 °C until analysis. After a 5 minutes centrifugation at 16,000 x g at 20 °C, the cleared supernatant was transferred to glass autosampler vials with 300 µl glass inserts (Chromatography Accessories Trott, Germany).

For the LC-HRMS analysis, 1 µl of the derivatized sample was injected onto a 100 x 2.1 mm HSS T3 UPLC column (Waters) with 1.8 µm particle size. The gradient elution ran in a binary Vanquish Horizon LC system (Thermo Fisher Scientific) with a flow rate of 400 µl/min using a binary buffer system consisting of buffer A :10 mM ammonium formate (Sigma), 0.15% [v/v] formic acid (Sigma) in LC-MS-grade water (Optima-Grade, Thermo Fisher Scientific) and buffer B consisted solely of acetonitrile (Optima-grade, Thermo Fisher-Scientific). The column temperature was set to 40 °C, while the LC gradient was: 0% B at 0 minutes, 0-15% B 0-4.1 minutes; 15-17% B 4.1 – 4.5 minutes; 17-55% B 4.5-11 minutes; 55-70% B 11 – 11.5 minutes, 70-100% B 11.5 – 13 minutes; B 100% 13 - 14 minutes; 100-0% B 14 -14.1 minutes; 0% B 14.1-19 minutes; 0% B. The mass spectrometer (Q-Exactive Plus, Thermo Fisher Scientific) was operating in positive ionization mode recording the mass range m/z 100-1000. The heated ESI source settings of the mass spectrometer were set to: Spray voltage 3.5 kV, capillary temperature 300 °C, sheath gas flow 60 AU, aux gas flow 20 AU at a temperature of 330 °C and the sweep gas to 2 AU. The RF-lens was set to a value of 60.

The identity of each compound was validated by authentic reference compounds, which were run before and after every sequence. Peak areas of [M + nBz + H]+ ions and isotopomers were extracted using a mass accuracy (<5 ppm) and a retention time tolerance of <0.05 minutes.

### Anion-Exchange Chromatography Mass Spectrometry (AEX-MS) for the analysis of anionic metabolites

Extracted metabolites were re-suspended in 200 µl of Optima UPLC/MS grade water (Thermo Fisher Scientific). After 15 minutes incubation on a thermomixer at 4 °C and a 5 minutes centrifugation at 16,000 x g at 4 °C, 100 µl of the cleared supernatant were transferred to polypropylene autosampler vials (Chromatography Accessories Trott).

The samples were analyzed using a Dionex ion chromatography system (Integrion, Thermo Fisher Scientific) as described previously (Schwaiger et al., 2017). In brief, 5 µl of polar metabolite extract was injected in full loop mode using an overfill factor of 1, onto a Dionex IonPac AS11-HC column (2 mm × 250 mm, 4 μm particle size, Thermo Fisher Scientific) equipped with a Dionex IonPac AG11-HC guard column (2 mm × 50 mm, 4 μm, Thermo Fisher Scientific). The column temperature was held at 30 °C, while the auto sampler was set to 6 °C. A potassium hydroxide gradient was generated using a potassium hydroxide cartridge (Eluent Generator, Thermo Scientific), which was supplied with deionized water. The metabolite separation was carried at a flow rate of 380 µl/min, applying the following gradient conditions: 0-minutes, 10 mM KOH; 3-12 minutes, 10−50 mM KOH; 12-19 minutes, 50-100 mM KOH, 19-21 minutes, 100 mM KOH, 21-22 minutes, 100-10 mM KOH. The column was re-equilibrated at 10 mM for 8 minutes.

For the analysis of metabolic pool sizes the eluting compounds were detected in negative ion mode using full scan measurements in the mass range m/z 50 – 750 on a Q-Exactive HF high resolution MS (Thermo Fisher Scientific). The heated ESI source settings of the mass spectrometer were: Spray voltage 3.2 kV, capillary temperature was set to 275 °C, sheath gas flow 70 AU, aux gas flow 15 AU at a temperature of 350 °C and a sweep gas flow of 0 AU. The S-lens was set to a value of 50.

The identity of each compound was validated by authentic reference compounds, which were measured at the beginning and the end of the sequence.

The LC/IC-MS data analysis was performed using the open source software El Maven 0.12.0 (Agrawal et al., 2019). For this purpose, Thermo raw mass spectra files were converted to mzML format using MSConvert 3.0.22060 (Chambers et al., 2012) (Proteowizard). Molecular ion and the corresponding isotopologue mass peaks of every required compound were extracted and integrated using the underlying algorithm within El Maven, only in rare cases isotopologue mass peaks were manually re-integrated. Extracted ion chromatograms were generated with a mass accuracy of <5 ppm and a retention time (RT) tolerance of <0.05 minutes as compared to the independently measured reference compounds. Afterwards, Isotopomer distribution was corrected for natural abundant isotopes with the IsoCorrectoR (Heinrich et al., 2018) package in R. If no independent 12C experiments were carried out, where the pool size is determined from the obtained peak area of the 12C monoisotopologue, the pool size determination was carried out by summing up the peak areas of all detectable isotopologues per compound. These areas were then normalized, as performed for un-traced 12C experiments, to the internal standards, which were added to the extraction buffer, followed by a normalization to the protein content or the cell number of the analyzed samples. The relative isotope distribution of each isotopologue was calculated from the proportion of the peak area of each isotopologue towards the sum of all detectable isotopologues. The ^13^C enrichment, namely the area attributed to ^13^C molecules traced in the detected isotopologues, was calculated by multiplying the peak area of each isotopologue with the proportion of the ^13^C and the ^12^C carbon number for the corresponding isotopologue (the ^12^C and ^13^C monoisotopologue areas were multiplied with 0 and 1 respectively). The obtained ^13^C area of each isotopologue are summed up, providing the peak area fraction associated to ^13^C atoms in the compound. Dividing this absolute ^13^C area by the summed area of all isotopologues provides the relative ^13^C enrichment factor.

## Supplementary Figure Text

**Supplementary Figure 1: Representative oxygen levels upon the addition of SB**

Representative oxygen level changes in control and 3 mM SB-treated fly heads (A), control and 15 mM SB-treated fly heads (B), control and 30 mM SB-treated fly heads (C). Each measurement lasted 2 minutes and consisted of 10 sub-measurements (ticks). The arrowhead indicates the addition of SB or buffer (control). The last measurement before (3) and all measurements after injection (4-18) are shown.

**Supplementary Figure 2: The protein levels of respiratory complex are not affected by the addition of SB after 20 minutes**

Levels of key mitochondrial proteins were not significantly altered after 20 minutes between control (orange, n=4) and 15 mM SB (green, n=4) group. Genes encoding proteins are as follows: (A) Complex I: CG6343 (ND-42), CG2286 (ND-75), (B) Complex II: CG17246 (SdhA), CG6666 (SdhC), (C) Complex III: CG8764 (Ox), CG7361 (Rieske), and (D) Complex IV: CG14724 (COX5A), CG9603 (COX7A). Error bars indicate ±S.E.M in all graphs. Student’s t-test was performed and detailed statistical information is presented in Supplementary Table 3.

**Supplementary Figure 3: The protein levels of key metabolic enzymes are not affected by the addition of SB after 20 minutes**

Levels of key mitochondrial proteins related to TCA cycle were not significantly altered after 20 minutes between control (orange, n=4) and 15 mM SB (green, n=4) group. Genes encoding proteins are as follows: (A) CG3861 (Citrate synthase), (B) CG9244 (Aconitate hydratase), (C) CG7176 (Isocitrate dehydrogenase), (D) CG1544 (2-oxoglutarate dehydrogenase), (E) CG5214 (Dihydrolipoyllysine-residue succinyltransferase), (F) CG7430 (Dihydrolipoyl dehydrogenase), (G) CG11963 (Succinyl-CoA ligase), (H) CG4094 (Fumarate hydratase), (I) CG5362 (Malate dehydrogenase). Error bars indicate ±S.E.M in all graphs. Student’s t-test was performed and detailed statistical information is presented in Supplementary Table 3.

**Supplementary Figure 4: Mass Isotopomer Distribution (MID) of butyric derivatives**

Analysis of Mass Isotopomer Distribution (MID) of butyric acid (A), butyryl-CoA (B), 3-hydroxybutyryl-CoA (C), and acetyl-CoA (D) ^13^C_4_-sodium butyrate treatment to determine its incorporation at different time points by mass spectrometry. Error bars indicate ±S.E.M in all graphs. n=6 for all measured metabolites and time points. M0 unlabeled mass of isotope, M+n native metabolite mass + number of isotopically labeled carbons. Detailed statistical analysis was performed using one-way ANOVA, Tukeýs post hoc test, results are presented in Supplementary Table 7.

**Supplementary Figure 5: Enrichment analysis of key metabolites in response to ^13^C_4_-labeled SB treatment**

(A-M) Enrichment analysis of key metabolites in response to ^13^C_4_-labeled SB treatment in fly heads at different time points (red, 0 min; blue, 20 min; green, 60 min). n=6 for all measured metabolites and time points, except for: succinyl-CoA 60 min n=4. Error bars indicate ±S.E.M in all graphs. One-way ANOVA, Tukeýs post hoc test, ns p>0.05; * p≤0.05; ** p≤0.01; *** p≤0.001; **** p≤0.0001. Detailed statistical information is presented in Supplementary Table 8.

**Supplementary Figure 6: Isotopomer Distribution of acetyl-CoA**

Analysis of Mass Isotopomer Distribution (MID) of acetyl-CoA (A) ^13^C_4_-sodium butyrate treatment to determine its incorporation at different time points by mass spectrometry. Error bars indicate ±S.E.M. n=6 for measured metabolite and time points. M0 unlabeled mass of isotope, M+n native metabolite mass + number of isotopically labeled carbons. Detailed statistical analysis was performed using one-way ANOVA, Tukeýs post hoc test, results are presented in Supplementary Table 7.

**Supplementary Figure 7: ^13^C labeling is not detected in amino acids after 20 minutes of labeled SB treatment** Analysis of Mass Isotopomer Distribution (MID)mass isotopomer distribution of representative metabolites: A) Arginine, B) Aspartate, C) Glutamate, D) Proline. Error bars indicate S.E.M, n=6 for measured metabolites and time points. M0 unlabeled mass of isotope, M+n native metabolite mass + number of isotopically labeled carbons.

**Supplementary Figure 8: Mass spectrometry analysis of 15 mM SB in the supernatant over time**

Supernatant samples were collected at three different time points (0 min, 20 min and 60 min) and analyzed by mass spectrometry for SB. The supernatant was collected from wells without fly heads (light green) and with fly heads (dark green) inside. 0 min, no fly heads n=3; 20 min, no fly heads n=4; 60 min, no fly heads n=4; 20 min, fly heads n=3; 60 min, fly heads n=3. Error bars indicate ±S.E.M in all groups. Two-way ANOVA, Tukeýs post hoc test, ns p>0.05. results are presented in Supplementary Table 9.

## Supplementary Tables

**Supplementary Table 1:**
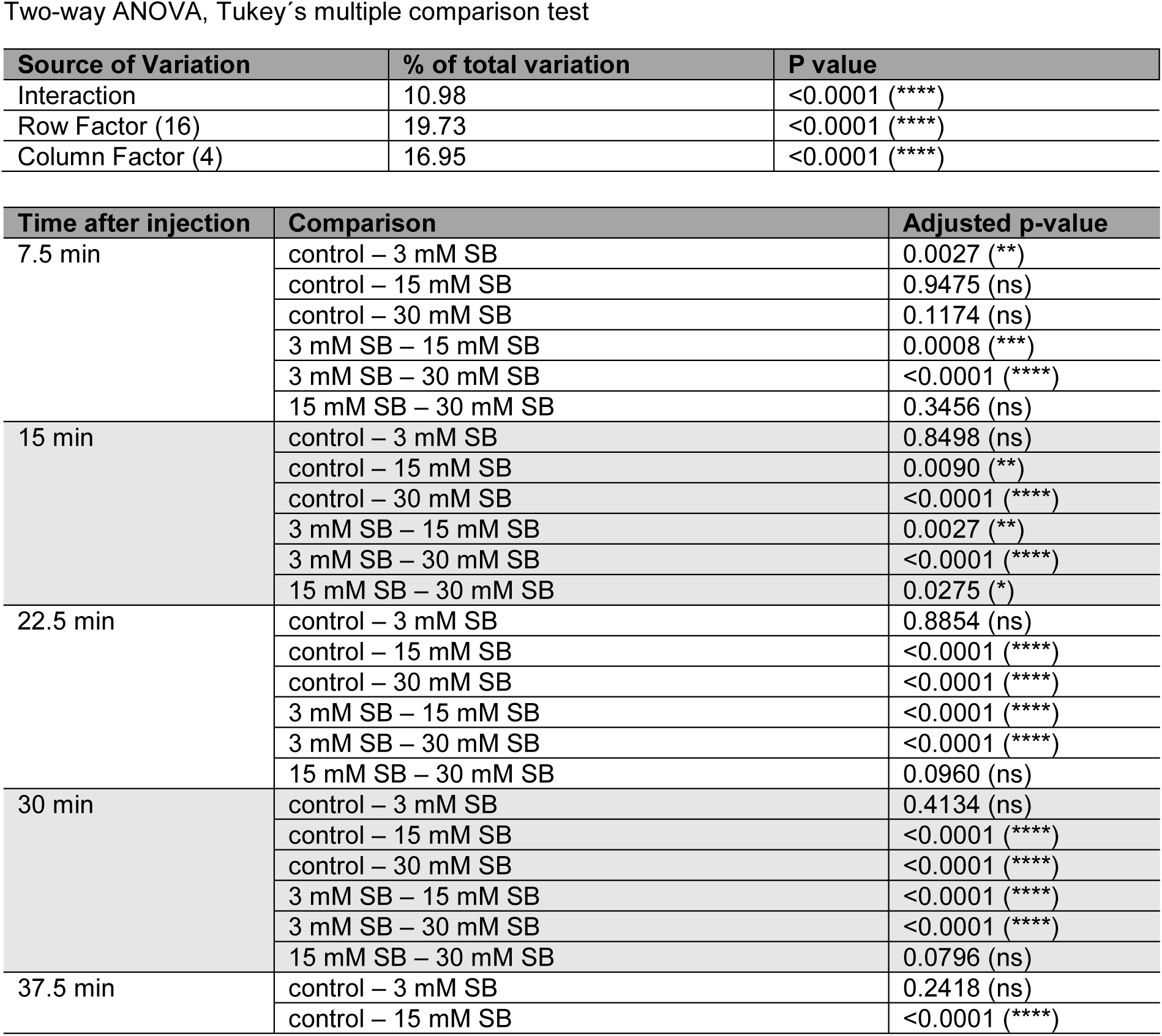
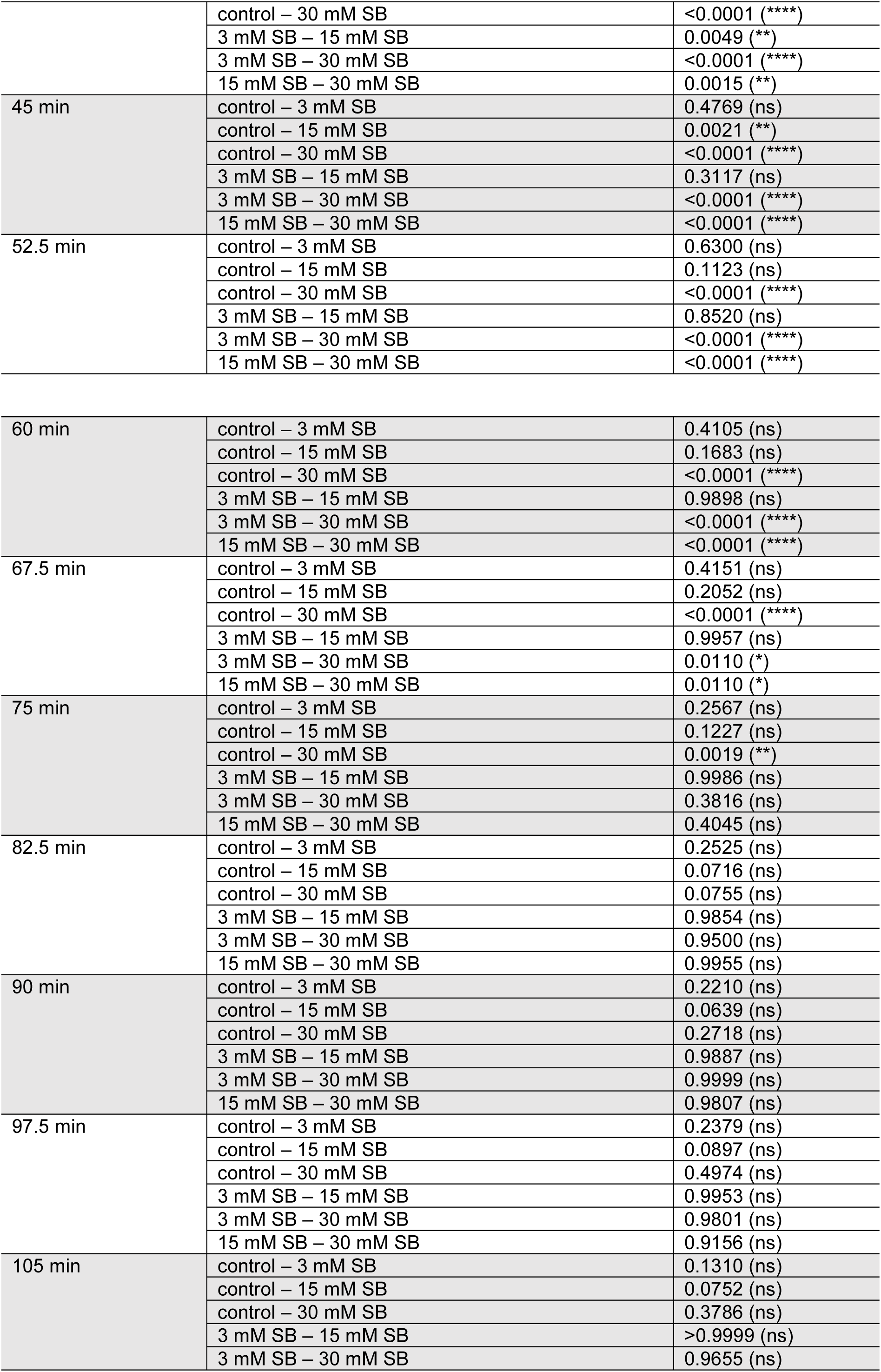
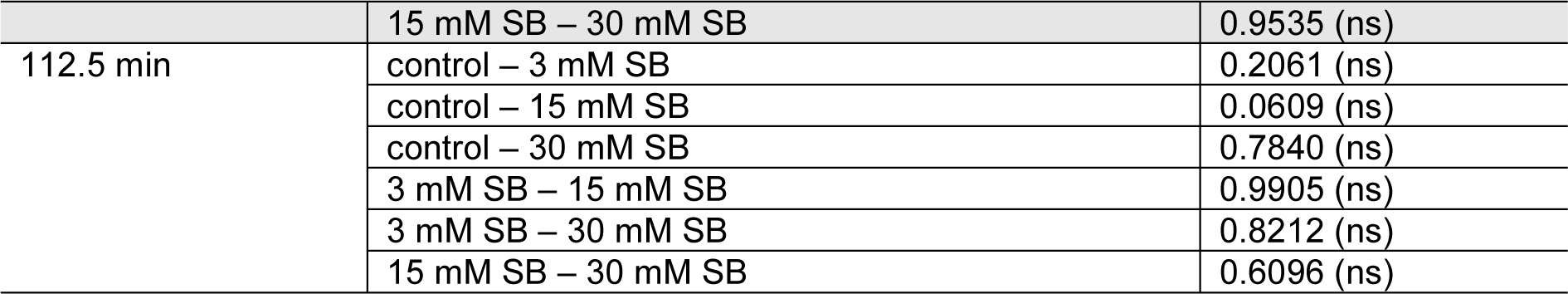
To Figure 1, A

**Supplementary Table 2:** To Figure 1, B, C:

| Figure | Condition | Mean ± SEM | n | CI 95% of mean |
| --- | --- | --- | --- | --- |
| 1, B | control | -1.413 ± 0.3346 | 28 | -2.099 – -0.7260 |
| | 3 mM SB | -1.062 ± 0.3598 | 16 | -1.829 – -0.2954 |
| | 15 mM SB | 2.066 ± 0.8069 | 22 | 0.3875 – 3.744 |
| | 30 mM SB | 4.463 ± 0.8803 | 15 | 2.575 – 6.352 |
| 1, C | control | -2.830 ± 0.4356 | 28 | -3.724 – -1.936 |
| | 3 mM SB | -1.647 ± 0.2967 | 16 | -2.279 – -1.015 |
| | 15 mM SB | -1.241 ± 0.3909 | 22 | -2.054 – -0.4280 |
| | 30 mM SB | 1.671 ± 0.3987 | 15 | 0.8161 – 2.526 |
| Figure | Comparison | Adjusted p-value | One-way ANOVA, Tukey's multiple comparison test |  |
| 1, B | control – 3 mM SB | 0.9770 (ns) | P value < 0.0001; R squared 0.4250 |  |
|  | control – 15 mM SB | 0.0002 (***) |  |  |
|  | control – 30 mM SB | < 0.0001 (****) |  |  |
|  | 3 mM SB – 15 mM SB | 0.0046 (**) |  |  |
|  | 3 mM SB – 30 mM SB | < 0.0001 (****) |  |  |
|  | 15 mM SB – 30 mM SB | 0.0519 (ns) |  |  |
| 1, C | control – 3 mM SB | 0.1892 (ns) | P value < 0.0001; R squared 0.4267 |  |
|  | control – 15 mM SB | 0.0194 (*) |  |  |
|  | control – 30 mM SB | < 0.0001 (****) |  |  |
|  | 3 mM SB – 15 mM SB | 0.9111 (ns) |  |  |
|  | 3 mM SB – 30 mM SB | < 0.0001 (****) |  |  |
|  | 15 mM SB – 30 mM SB | < 0.0001 (****) |  |  |

**Supplementary Table 3:** To Supplementary Figure 2, 3

**Proteome**
| Protein ID | CG number | Control | Mean ± SEM | SB | Mean ± SEM | Student t-test adjusted p-Value |
| --- | --- | --- | --- | --- | --- | --- |
| P91929<br>(NADH dehydrogenase [ubiquinone] 1 alpha subcomplex subunit 10) | CG6343<br>(ND-42) | 9.4224 | 9.3960 ± 0.0223 | 9.3791 | 9.4083 ± 0.0113 | 0.9698 |
|  |  | 9.3975 |  | 9.4088 |  |  |
|  |  | 9.3329 |  | 9.4344 |  |  |
|  |  | 9.4320 |  | 9.4111 |  |  |
| Q94511 |  | 9.8504 |  | 9.8714 |  | 0.9449 |
| (NADH-ubiquinone<br>oxidoreductase 75 kDa<br>subunit) | CG2286<br>(ND-75) | 9.8497 | 9.8633<br>±<br>0.0082 | 9.8457 | 9.8756<br>±<br>0.0132 |  |
|  |  | 9.8838 |  | 9.9099 |  |  |
|  |  | 9.8691 |  | 9.8752 |  |  |
| Q94523<br>(Succinate dehydrogenase<br>[ubiquinone] flavoprotein<br>subunit) | CG17246<br>(SdhA) | 10.0389 | 10.0286<br>±<br>0.0064 | 10.0535 | 10.0404<br>±<br>0.0070 | 0.9449 |
|  |  | 10.0341 |  | 10.0207 |  |  |
|  |  | 10.0099 |  | 10.0415 |  |  |
|  |  | 10.0316 |  | 10.0457 |  |  |
| Q9VGS3<br>(Succinate dehydrogenase<br>cytochrome b560 subunit) | CG6666<br>(SdhC) | 8.7732 | 8.8062<br>±<br>0.0155 | 8.7039 | 8.8031<br>±<br>0.0396 | 0.9983 |
|  |  | 8.8249 |  | 8.7884 |  |  |
|  |  | 8.8391 |  | 8.8264 |  |  |
|  |  | 8.7876 |  | 8.8939 |  |  |
| Q9XY35<br>(Cytochrome b-c1 complex subunit 9) | CG8764<br>(ox) | 9.1050 | 9.0880<br>±<br>0.0184 | 9.0835 | 9.1133<br>±<br>0.0152 | 0.9449 |
|  |  | 9.1014 |  | 9.1305 |  |  |
|  |  | 9.0334 |  | 9.0922 |  |  |
|  |  | 9.1123 |  | 9.1472 |  |  |
| Q9VQ29<br>(Cytochrome b-c1 complex subunit Rieske) | CG7361<br>(RFeSP) | 9.6808 | 9.6663<br>±<br>0.0105 | 9.6726 | 9.6621<br>±<br>0.0143 | 0.9774 |
|  |  | 9.6350 |  | 9.6784 |  |  |
|  |  | 9.6762 |  | 9.6193 |  |  |
|  |  | 9.6730 |  | 9.6782 |  |  |
| Q94514<br>(Cytochrome c oxidase subunit 5A) | CG14724<br>(COX5A) | 9.9490 | 9.9225<br>±<br>0.0202 | 9.9412 | 9.9427<br>±<br>0.0054 | 0.9449 |
|  |  | 9.9320 |  | 9.9532 |  |  |
|  |  | 9.8628 |  | 9.9482 |  |  |
|  |  | 9.9461 |  | 9.9281 |  |  |
| Q9VHS2 (Cytochrome c oxidase subunit 7A) | CG9603<br>(COX7A) | 9.4119 | 9.4116<br>±<br>0.0069 | 9.4161 | 9.4198<br>±<br>0.0027 | 0.9449 |
|  |  | 9.3974 |  | 9.4171 |  |  |
|  |  | 9.4069 |  | 9.4179 |  |  |
|  |  | 9.4301 |  | 9.4279 |  |  |
| P35381 (ATP synthase subunit alpha) | CG3612<br>(blw) | 11.0295 | 11.0238<br>±<br>0.0028 | 11.0406 | 11.0249<br>±<br>0.0085 | 0.9956 |
|  |  | 11.0196 |  | 11.0227 |  |  |
|  |  | 11.0278 |  | 11.0343 |  |  |
|  |  | 11.0184 |  | 11.0019 |  |  |
| Q24251 (ATP synthase subunit d) | CG6030<br>(ATPsyn-d) | 9.9758 | 9.9332<br>±<br>0.0244 | 9.9362 | 9.9442<br>±<br>0.0080 | 0.9713 |
|  |  | 9.9355 |  | 9.9681 |  |  |
|  |  | 9.8642 |  | 9.9364 |  |  |
|  |  | 9.9573 |  | 9.9362 |  |  |
| Q9W401 (Probable citrate synthase) | CG3861<br>(kdn) | 10.5849 | 10.5707<br>±<br>0.0052 | 10.5991 | 10.5810<br>±<br>0.0074 | 0.9449 |
|  |  | 10.5688 |  | 10.5653 |  |  |
|  |  | 10.5688 |  | 10.5864 |  |  |
|  |  | 10.5601 |  | 10.5734 |  |  |
| Q9VIE8<br>(Aconitate hydratase) | CG9244<br>(mAcon1) | 10.6498 | 10.6520<br>±<br>0.0076 | 10.6935 | 10.6662<br>±<br>0.0095 | 0.9449 |
|  |  | 10.6729 |  | 10.6624 |  |  |
|  |  | 10.6488 |  | 10.6590 |  |  |
|  |  | 10.6365 |  | 10.6497 |  |  |
| Q7KUB1<br>(Isocitrate dehydrogenase [NADP]) | CG7176<br>(ldh) | 10.3528 | 10.3589<br>±<br>0.0040 | 10.3557 | 10.3568<br>±<br>0.0037 | 0.9713 |
|  |  | 10.3532 |  | 10.3574 |  |  |
|  |  | 10.3599 |  | 10.3479 |  |  |
|  |  | 10.3697 |  | 10.3661 |  |  |
| Q9VA02<br>(Probable 2-oxoglutarate dehydrogenase E1 component DHKTD1 homolog) | CG1544 | 8.8059 | 8.8110<br>±<br>0.0085 | 8.8028 | 8.8151<br>±<br>0.0094 | 0.9759 |
|  |  | 8.8036 |  | 8.7956 |  |  |
|  |  | 8.7985 |  | 8.8289 |  |  |
|  |  | 8.8361 |  | 8.8332 |  |  |
| Q9VGQ1<br>(Dihydrolipoyllysine-residue succinyltransferase component of 2-oxoglutarate dehydrogenase complex) | CG5214 | 9.7823 | 9.7797<br>±<br>0.0009 | 9.7817 | 9.7888<br>±<br>0.0060 | 0.9449 |
|  |  | 9.7791 |  | 9.7887 |  |  |
|  |  | 9.7785 |  | 9.8057 |  |  |
|  |  | 9.7788 |  | 9.7792 |  |  |
| Q9VVL7 (Dihydrolipoyl dehydrogenase) | CG7430 | 10.0387 | 10.0342<br>±<br>0.0029 | 10.0382 | 10.0384<br>±<br>0.0096 | 0.9713 |
|  |  | 10.0289 |  | 10.0391 |  |  |
|  |  | 10.0398 |  | 10.0147 |  |  |
|  |  | 10.0293 |  | 10.0616 |  |  |
| Q95U38<br>(Succinate--CoA ligase [ADP-forming] subunit beta) | CG11963<br>(skap) | 10.2413 | 10.2056<br>±<br>0.0197 | 10.2254 | 10.2150<br>±<br>0.0050 | 0.9713 |
|  |  | 10.2022 |  | 10.2132 |  |  |
|  |  | 10.1518 |  | 10.2020 |  |  |
|  |  | 10.2271 |  | 10.2196 |  |  |
| Q8IRQ5<br>(fumarate hydratase) | CG4094<br>(Fum1) | 9.8735 | 9.8947<br>±<br>0.0158 | 9.8645 | 9.8849<br>±<br>0.0108 | 0.9675 |
|  |  | 9.8653 |  | 9.8808 |  |  |
|  |  | 9.9345 |  | 9.9155 |  |  |
|  |  | 9.9054 |  | 9.8789 |  |  |
| Q9VKX2<br>(Malate dehydrogenase) | CG5362<br>(Mdh1) | 10.0215 | 10.0224<br>±<br>0.0050 | 10.0325 | 10.0263<br>±<br>0.0068 | 0.9698 |
|  |  | 10.0261 |  | 10.0249 |  |  |
|  |  | 10.0092 |  | 10.0080 |  |  |
|  |  | 10.0327 |  | 10.0400 |  |  |

**Supplementary Table 4:** To Figure 3, B

Two-way ANOVA, Tukey's multiple comparison test
| Source of Variation | % of total variation | P value |
| --- | --- | --- |
| Interaction | 5.057 | 0.3102 (ns) |
| Row Factor (16) | 9.491 | <0.0001 (****) |
| Column Factor (3) | 14.68 | <0.0001 (****) |

| Time after injection | Comparison | Adjusted p-value |
| --- | --- | --- |
| 7.5 min | control – 5 mM PP | 0.7662 (ns) |
|  | control – 25 mM PP | 0.0524 (ns) |
|  | 5 mM PP – 25 mM PP | 0.1515 (ns) |
| 15 min | control – 5 mM PP | 0.1123 (ns) |
|  | control – 25 mM PP | 0.0032 (**) |
|  | 5 mM PP – 25 mM PP | 0.1964 (ns) |
| 22.5 min | control – 5 mM PP | 0.0486 (*) |
|  | control – 25 mM PP | 0.0006 (***) |
|  | 5 mM PP – 25 mM PP | 0.1358 (ns) |
| 30 min | control – 5 mM PP | 0.2282 (ns) |
|  | control – 25 mM PP | 0.0001 (***) |
|  | 5 mM PP – 25 mM PP | 0.0089 (**) |
| 37.5 min | control – 5 mM PP | 0.4050 (ns) |
|  | control – 25 mM PP | 0.0002 (***) |
|  | 5 mM PP – 25 mM PP | 0.0050 (**) |
| 45 min | control – 5 mM PP | 0.4655 (ns) |
|  | control – 25 mM PP | 0.0003 (***) |
|  | 5 mM PP – 25 mM PP | 0.0047 (**) |
| 52.5 min | control – 5 mM PP | 0.4269 (ns) |
|  | control – 25 mM PP | 0.0003 (***) |
|  | 5 mM PP – 25 mM PP | 0.0061 (**) |
| 60 min | control – 5 mM PP | 0.4785 (ns) |
|  | control – 25 mM PP | 0.0013 (**) |
|  | 5 mM PP – 25 mM PP | 0.0185 (*) |
| 67.5 min | control – 5 mM PP | 0.4867 (ns) |
|  | control – 25 mM PP | 0.0137 (*) |
|  | 5 mM PP – 25 mM PP | 0.1159 (ns) |
| 75 min | control – 5 mM PP | 0.5303 (ns) |
|  | control – 25 mM PP | 0.1061 (ns) |
|  | 5 mM PP – 25 mM PP | 0.4464 (ns) |
| 82.5 min | control – 5 mM PP | 0.5928 (ns) |
|  | control – 25 mM PP | 0.2190 (ns) |
|  | 5 mM PP – 25 mM PP | 0.6312 (ns) |
| 90 min | control – 5 mM PP | 0.5929 (ns) |
|  | control – 25 mM PP | 0.3860 (ns) |
|  | 5 mM PP – 25 mM PP | 0.8486 (ns) |
| 97.5 min | control – 5 mM PP | 0.7765 (ns) |
|  | control – 25 mM PP | 0.6859 (ns) |
|  | 5 mM PP – 25 mM PP | 0.9558 (ns) |
| 105 min | control – 5 mM PP | 0.5832 (ns) |
|  | control – 25 mM PP | 0.6280 (ns) |
|  | 5 mM PP – 25 mM PP | 0.9930 (ns) |
| 112.5 min | control – 5 mM PP | 0.5671 (ns) |
|  | control – 25 mM PP | 0.6668 (ns) |
|  | 5 mM PP – 25 mM PP | 0.9996 (ns) |

**Supplementary Table 5:** To Figure 3, C, D

| Figure | Condition | Mean ± SEM | n | CI 95% of mean |
| --- | --- | --- | --- | --- |
| 3, C | control | 0.4647 ± 0.8318 | 11 | -1.389 – 2.318 |
| | 5 mM PP | 3.240 ± 1.357 | 15 | 0.3300 – 6.150 |
| | 25 mM PP | 8.008 ± 2.335 | 7 | 2.295 – 13.72 |
| 3, D | control | -0.5767 ± 0.9994 | 11 | -2.803 – 1.650 |
| | 5 mM PP | 1.346 ± 0.9356 | 15 | -0.6612 – 3.352 |
| | 25 mM PP | 5.639 ± 1.699 | 7 | 1.482 – 9.795 |
| Figure | Comparison | Adjusted p-value | One-way ANOVA, Tukey's multiple comparison test |  |
| 3, C | control – 5 mM PP | 0.3452 (ns) | P value 0.0438; R squared 0.1882 |  |
|  | control – 25 mM PP | 0.0346 (*) |  |  |
|  | 5 mM PP – 25 mM PP | 0.2831 (ns) |  |  |
| 3, D | control – 5 mM PP | 0.3751 (ns) | P value 0.0028; R squared 0.3235 |  |
|  | control – 25 mM PP | 0.0021 (**) |  |  |
|  | 5 mM PP – 25 mM PP | 0.0239 (*) |  |  |

**Supplementary Table 6:**
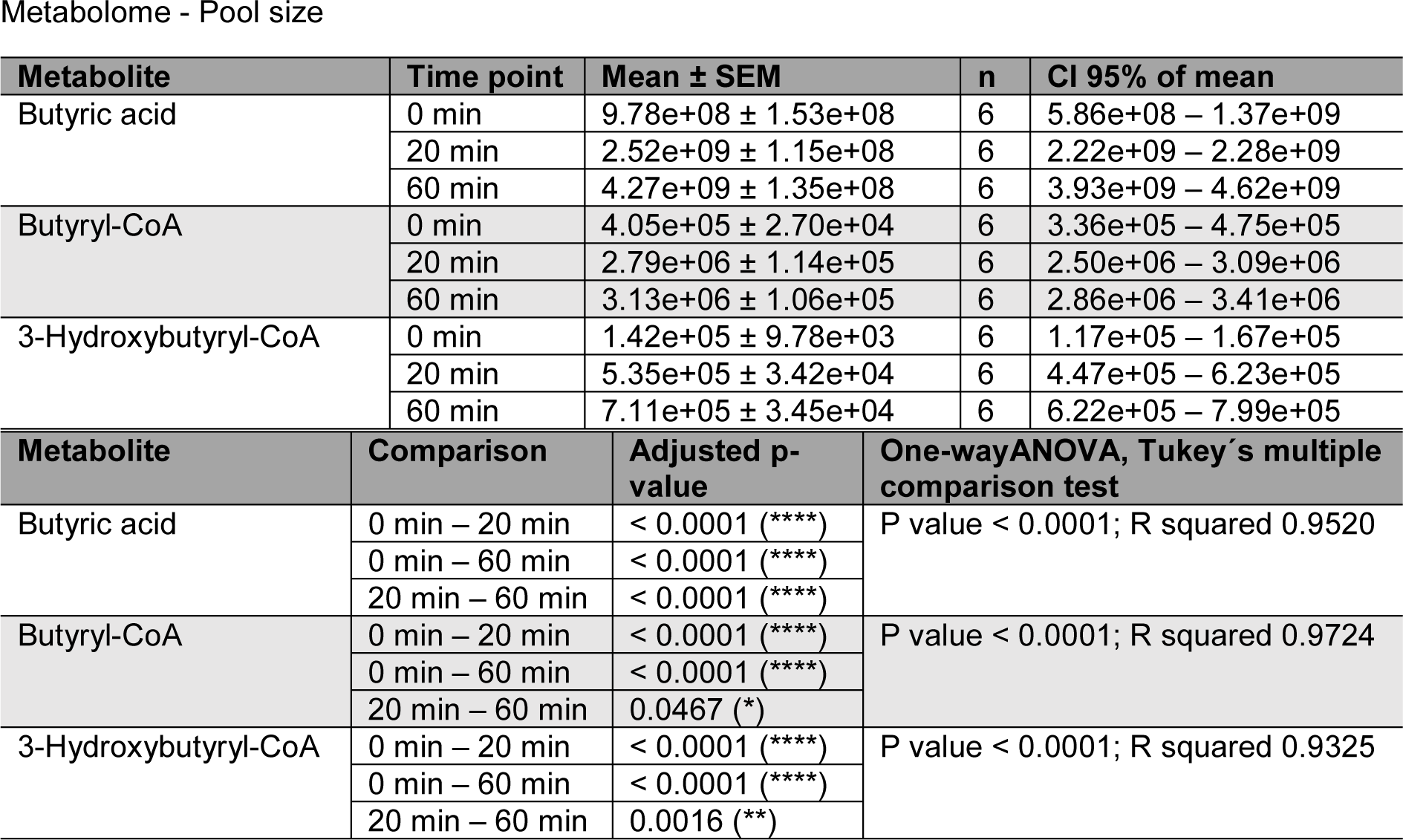
To Figure 4

**Supplementary Table 7:**
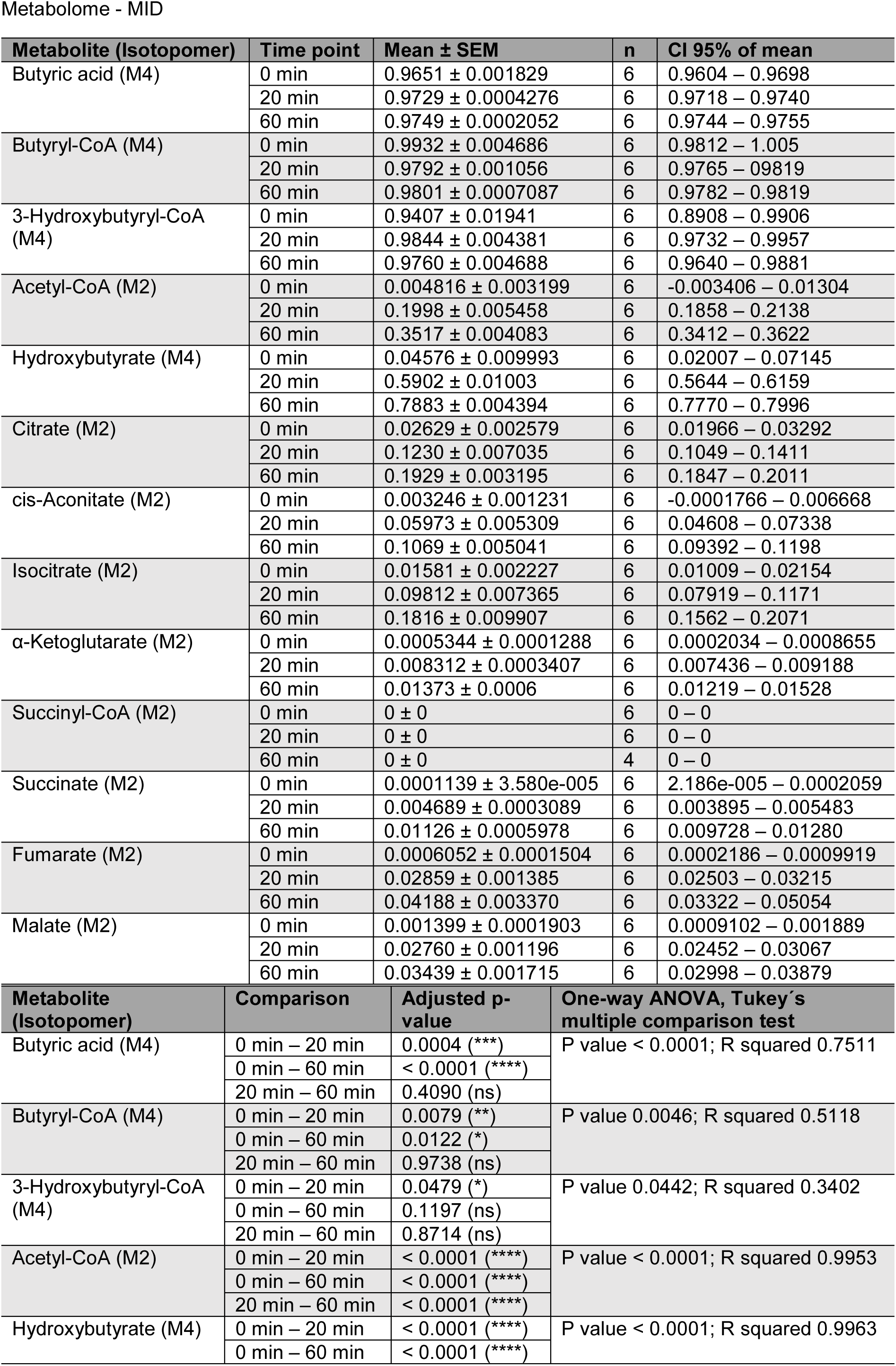
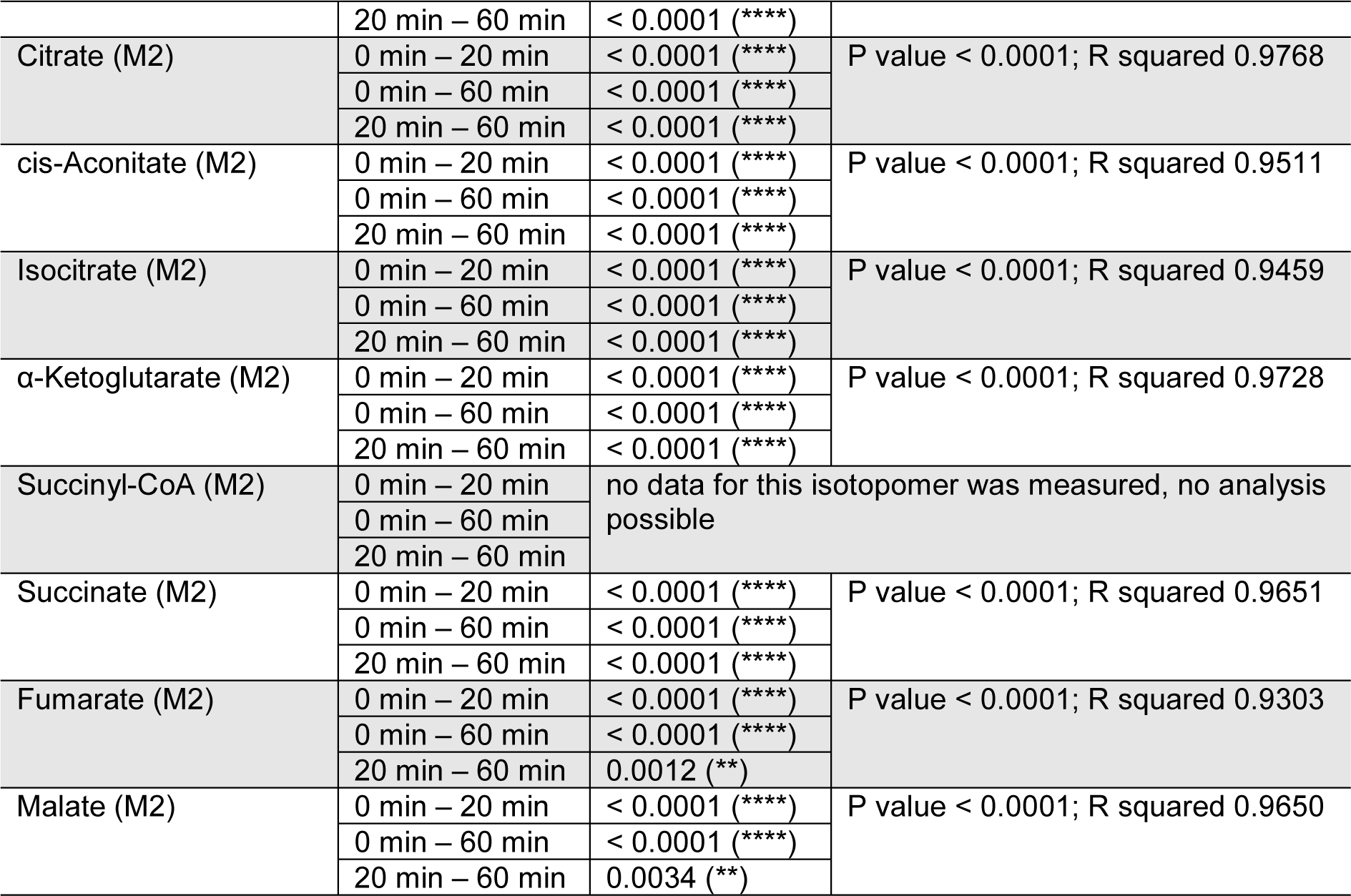
To Figure 5 and Supplementary Figure 4, 6

| Metabolite (Isotopomer) | Time point | Mean ± SEM | n | CI 95% of mean |
| --- | --- | --- | --- | --- |
| Butyric acid (M4) | 0 min | 0.9651 ± 0.001829 | 6 | 0.9604 – 0.9698 |
| | 20 min | 0.9729 ± 0.0004276 | 6 | 0.9718 – 0.9740 |
| | 60 min | 0.9749 ± 0.0002052 | 6 | 0.9744 – 0.9755 |
| Butyryl-CoA (M4) | 0 min | 0.9932 ± 0.004686 | 6 | 0.9812 – 1.005 |
| | 20 min | 0.9792 ± 0.001056 | 6 | 0.9765 – 0.9819 |
| | 60 min | 0.9801 ± 0.0007087 | 6 | 0.9782 – 0.9819 |
| 3-Hydroxybutyryl-CoA (M4) | 0 min | 0.9407 ± 0.01941 | 6 | 0.8908 – 0.9906 |
| | 20 min | 0.9844 ± 0.004381 | 6 | 0.9732 – 0.9957 |
| | 60 min | 0.9760 ± 0.004688 | 6 | 0.9640 – 0.9881 |
| Acetyl-CoA (M2) | 0 min | 0.004816 ± 0.003199 | 6 | -0.003406 – 0.01304 |
| | 20 min | 0.1998 ± 0.005458 | 6 | 0.1858 – 0.2138 |
| | 60 min | 0.3517 ± 0.004083 | 6 | 0.3412 – 0.3622 |
| Hydroxybutyrate (M4) | 0 min | 0.04576 ± 0.009993 | 6 | 0.02007 – 0.07145 |
| | 20 min | 0.5902 ± 0.01003 | 6 | 0.5644 – 0.6159 |
| | 60 min | 0.7883 ± 0.004394 | 6 | 0.7770 – 0.7996 |
| Citrate (M2) | 0 min | 0.02629 ± 0.002579 | 6 | 0.01966 – 0.03292 |
| | 20 min | 0.1230 ± 0.007035 | 6 | 0.1049 – 0.1411 |
| | 60 min | 0.1929 ± 0.003195 | 6 | 0.1847 – 0.2011 |
| cis-Aconitate (M2) | 0 min | 0.003246 ± 0.001231 | 6 | -0.0001766 – 0.006668 |
| | 20 min | 0.05973 ± 0.005309 | 6 | 0.04608 – 0.07338 |
| | 60 min | 0.1069 ± 0.005041 | 6 | 0.09392 – 0.1198 |
| Isocitrate (M2) | 0 min | 0.01581 ± 0.002227 | 6 | 0.01009 – 0.02154 |
| | 20 min | 0.09812 ± 0.007365 | 6 | 0.07919 – 0.1171 |
| | 60 min | 0.1816 ± 0.009907 | 6 | 0.1562 – 0.2071 |
| α-Ketoglutarate (M2) | 0 min | 0.0005344 ± 0.0001288 | 6 | 0.0002034 – 0.0008655 |
| | 20 min | 0.008312 ± 0.0003407 | 6 | 0.007436 – 0.009188 |
| | 60 min | 0.01373 ± 0.0006 | 6 | 0.01219 – 0.01528 |
| Succinyl-CoA (M2) | 0 min | 0 ± 0 | 6 | 0 – 0 |
| | 20 min | 0 ± 0 | 6 | 0 – 0 |
| | 60 min | 0 ± 0 | 4 | 0 – 0 |
| Succinate (M2) | 0 min | 0.0001139 ± 3.580e-005 | 6 | 2.186e-005 – 0.0002059 |
| | 20 min | 0.004689 ± 0.0003089 | 6 | 0.003895 – 0.005483 |
| | 60 min | 0.01126 ± 0.0005978 | 6 | 0.009728 – 0.01280 |
| Fumarate (M2) | 0 min | 0.0006052 ± 0.0001504 | 6 | 0.0002186 – 0.0009919 |
| | 20 min | 0.02859 ± 0.001385 | 6 | 0.02503 – 0.03215 |
| | 60 min | 0.04188 ± 0.003370 | 6 | 0.03322 – 0.05054 |
| Malate (M2) | 0 min | 0.001399 ± 0.0001903 | 6 | 0.0009102 – 0.001889 |
| | 20 min | 0.02760 ± 0.001196 | 6 | 0.02452 – 0.03067 |
| | 60 min | 0.03439 ± 0.001715 | 6 | 0.02998 – 0.03879 |
| Metabolite (Isotopomer) | Comparison | Adjusted p-value | One-way ANOVA, Tukey's multiple comparison test |  |
| Butyric acid (M4) | 0 min – 20 min | 0.0004 (***) | P value < 0.0001; R squared 0.7511 |  |
|  | 0 min – 60 min | < 0.0001 (****) |  |  |
|  | 20 min – 60 min | 0.4090 (ns) |  |  |
| Butyryl-CoA (M4) | 0 min – 20 min | 0.0079 (**) | P value 0.0046; R squared 0.5118 |  |
|  | 0 min – 60 min | 0.0122 (*) |  |  |
|  | 20 min – 60 min | 0.9738 (ns) |  |  |
| 3-Hydroxybutyryl-CoA (M4) | 0 min – 20 min | 0.0479 (*) | P value 0.0442; R squared 0.3402 |  |
|  | 0 min – 60 min | 0.1197 (ns) |  |  |
|  | 20 min – 60 min | 0.8714 (ns) |  |  |
| Acetyl-CoA (M2) | 0 min – 20 min | < 0.0001 (****) | P value < 0.0001; R squared 0.9953 |  |
|  | 0 min – 60 min | < 0.0001 (****) |  |  |
|  | 20 min – 60 min | < 0.0001 (****) |  |  |
| Hydroxybutyrate (M4) | 0 min – 20 min | < 0.0001 (****) | P value < 0.0001; R squared 0.9963 |  |
|  | 0 min – 60 min | < 0.0001 (****) |  |  |

|  |  |  |  |
| --- | --- | --- | --- |
|  | 20 min – 60 min | < 0.0001 (****) |  |
| Citrate (M2) | 0 min – 20 min | < 0.0001 (****) | P value < 0.0001; R squared 0.9768 |
|  | 0 min – 60 min | < 0.0001 (****) |  |
|  | 20 min – 60 min | < 0.0001 (****) |  |
| cis-Aconitate (M2) | 0 min – 20 min | < 0.0001 (****) | P value < 0.0001; R squared 0.9511 |
|  | 0 min – 60 min | < 0.0001 (****) |  |
|  | 20 min – 60 min | < 0.0001 (****) |  |
| Isocitrate (M2) | 0 min – 20 min | < 0.0001 (****) | P value < 0.0001; R squared 0.9459 |
|  | 0 min – 60 min | < 0.0001 (****) |  |
|  | 20 min – 60 min | < 0.0001 (****) |  |
| α-Ketoglutarate (M2) | 0 min – 20 min | < 0.0001 (****) | P value < 0.0001; R squared 0.9728 |
|  | 0 min – 60 min | < 0.0001 (****) |  |
|  | 20 min – 60 min | < 0.0001 (****) |  |
| Succinyl-CoA (M2) | 0 min – 20 min | no data for this isotopomer was measured, no analysis possible |  |
|  | 0 min – 60 min |  |  |
|  | 20 min – 60 min |  |  |
| Succinate (M2) | 0 min – 20 min | < 0.0001 (****) | P value < 0.0001; R squared 0.9651 |
|  | 0 min – 60 min | < 0.0001 (****) |  |
|  | 20 min – 60 min | < 0.0001 (****) |  |
| Fumarate (M2) | 0 min – 20 min | < 0.0001 (****) | P value < 0.0001; R squared 0.9303 |
|  | 0 min – 60 min | < 0.0001 (****) |  |
|  | 20 min – 60 min | 0.0012 (**) |  |
| Malate (M2) | 0 min – 20 min | < 0.0001 (****) | P value < 0.0001; R squared 0.9650 |
|  | 0 min – 60 min | < 0.0001 (****) |  |
|  | 20 min – 60 min | 0.0034 (**) |  |

**Supplementary Table 8:** To Supplementary Figure 5

| Metabolite | Time point | Mean ± SEM | n | CI 95% of mean |
| --- | --- | --- | --- | --- |
| Butyric acid | 0 min | 97.88 ± 0.1939 | 6 | 97.38 – 98.38 |
|  | 20 min | 98.74 ± 0.04604 | 6 | 98.62 – 98.85 |
|  | 60 min | 98.95 ± 0.02124 | 6 | 98.90 – 99.01 |
| Butyryl-CoA | 0 min | 99.32 ± 0.4686 | 6 | 98.12 – 100.5 |
|  | 20 min | 99.34 ± 0.06858 | 6 | 99.17 – 99.52 |
|  | 60 min | 99.50 ± 0.01772 | 6 | 99.46 – 99.55 |
| 3-Hydroxybutyryl-CoA | 0 min | 94.07 ± 1.941 | 6 | 89.08 – 99.06 |
|  | 20 min | 99.34 ± 0.1815 | 6 | 98.88 – 99.81 |
|  | 60 min | 99.07 ± 0.1847 | 6 | 98.59 – 99.54 |
| Acetyl-CoA | 0 min | 0.4816 ± 0.3199 | 6 | -0.3406 – 1.304 |
|  | 20 min | 19.98 ± 0.5458 | 6 | 18.58 – 21.38 |
|  | 60 min | 35.17 ± 0.4083 | 6 | 34.12 – 36.22 |
| Hydroxybutyrate | 0 min | 4.576 ± 0.9993 | 6 | 2.007 – 7.145 |
|  | 20 min | 59.55 ± 1.030 | 6 | 56.90 – 62.20 |
|  | 60 min | 80.04 ± 0.4568 | 6 | 78.87 – 81.22 |
| Citrate | 0 min | 0.9750 ± 0.09758 | 6 | 0.7241 – 1.226 |
|  | 20 min | 5.633 ± 0.3063 | 6 | 4.845 – 6.420 |
|  | 60 min | 10.76 ± 0.2363 | 6 | 10.15 – 11.36 |
| cis-Aconitate | 0 min | 0.1082 ± 0.04438 | 6 | -0.005886 – 0.2223 |
|  | 20 min | 2.025 ± 0.2010 | 6 | 1.509 – 2.542 |
|  | 60 min | 3.698 ± 0.1842 | 6 | 3.225 – 4.171 |
| Isocitrate | 0 min | 0.5272 ± 0.07423 | 6 | 0.3363 – 0.7180 |
|  | 20 min | 3.416 ± 0.2555 | 6 | 2.759 – 4.073 |
|  | 60 min | 7.765 ± 0.5219 | 6 | 6.423 – 9.106 |

| α-Ketoglutarate | 0 min | $0.2138 \pm 0.005152$ | 6 | $0.008134 - 0.03462$ |
| --- | --- | --- | --- | --- |
| | 20 min | $0.4388 \pm 0.02331$ | 6 | $0.3789 - 0.4987$ |
| | 60 min | $1.026 \pm 0.06081$ | 6 | $0.8697 - 1.182$ |
| Succinyl-CoA | 0 min | $0.05686 \pm 0.05686$ | 6 | $-0.08930 - 0.2030$ |
| | 20 min | $0 \pm 0$ | 6 | $0 - 0$ |
| | 60 min | $0 \pm 0$ | 4 | $0 - 0$ |
| Succinate | 0 min | $0.04172 \pm 0.008589$ | 6 | $0.01964 - 0.06380$ |
| | 20 min | $0.3410 \pm 0.01696$ | 6 | $0.2974 - 0.3846$ |
| | 60 min | $0.8723 \pm 0.04888$ | 6 | $0.7467 - 0.9980$ |
| Fumarate | 0 min | $0.03026 \pm 0.007521$ | 6 | $0.01093 - 0.04959$ |
| | 20 min | $1.956 \pm 0.1039$ | 6 | $1.689 - 2.223$ |
| | 60 min | $3.984 \pm 0.3929$ | 6 | $2.974 - 4.994$ |
| Malate | 0 min | $0.07637 \pm 0.008878$ | 6 | $0.05355 - 0.09920$ |
| | 20 min | $2.013 \pm 0.09125$ | 6 | $1.778 - 2.247$ |
| | 60 min | $2.983 \pm 0.1951$ | 6 | $2.482 - 3.485$ |
| Metabolite | Comparison | Adjusted p-value | One-way ANOVA, Tukey's multiple comparison test |  |
| Butyric acid | 0 min – 20 min | 0.0003 (***) | P value < 0.0001; R squared 0.7633 |  |
|  | 0 min – 60 min | < 0.0001 (****) |  |  |
|  | 20 min – 60 min | 0.4047 (ns) |  |  |
| Butyryl-CoA | 0 min – 20 min | 0.9984 (ns) | P value 0.8794; R squared 0.01699 |  |
|  | 0 min – 60 min | 0.8885 (ns) |  |  |
|  | 20 min – 60 min | 0.9113 (ns) |  |  |
| 3-Hydroxybutyryl-CoA | 0 min – 20 min | 0.0127 (*) | P value 0.0075; R squared 0.4789 |  |
|  | 0 min – 60 min | 0.0179 (*) |  |  |
|  | 20 min – 60 min | 0.9837 (ns) |  |  |
| Acetyl-CoA | 0 min – 20 min | < 0.0001 (****) | P value < 0.0001; R squared 0.9953 |  |
|  | 0 min – 60 min | < 0.0001 (****) |  |  |
|  | 20 min – 60 min | < 0.0001 (****) |  |  |
| Hydroxybutyrate | 0 min – 20 min | < 0.0001 (****) | P value < 0.0001; R squared 0.9963 |  |
|  | 0 min – 60 min | < 0.0001 (****) |  |  |
|  | 20 min – 60 min | < 0.0001 (****) |  |  |
| Citrate | 0 min – 20 min | < 0.0001 (****) | P value < 0.0001; R squared 0.9836 |  |
|  | 0 min – 60 min | < 0.0001 (****) |  |  |
|  | 20 min – 60 min | < 0.0001 (****) |  |  |
| cis-Aconitate | 0 min – 20 min | < 0.0001 (****) | P value < 0.0001; R squared 0.9442 |  |
|  | 0 min – 60 min | < 0.0001 (****) |  |  |
|  | 20 min – 60 min | < 0.0001 (****) |  |  |
| Isocitrate | 0 min – 20 min | < 0.0001 (****) | P value < 0.0001; R squared 0.9393 |  |
|  | 0 min – 60 min | < 0.0001 (****) |  |  |
|  | 20 min – 60 min | < 0.0001 (****) |  |  |
| α-Ketoglutarate | 0 min – 20 min | < 0.0001 (****) | P value < 0.0001; R squared 0.9598 | |
|  | 0 min – 60 min | < 0.0001 (****) |  |  |
|  | 20 min – 60 min | < 0.0001 (****) |  |  |
| Succinyl-CoA | 0 min – 20 min | 0.5076 (ns) | P value 0.4651; R squared 0.1111 |  |
|  | 0 min – 60 min | 0.5779 (ns) |  |  |
|  | 20 min – 60 min | > 0.9999 (ns) |  |  |
| Succinate | 0 min – 20 min | < 0.0001 (****) | P value < 0.0001; R squared 0.9626 |  |
|  | 0 min – 60 min | < 0.0001 (****) |  |  |
|  | 20 min – 60 min | < 0.0001 (****) |  |  |
| Fumarate | 0 min – 20 min | < 0.0001 (****) | P value < 0.0001; R squared 0.9044 |  |
|  | 0 min – 60 min | < 0.0001 (****) |  |  |
|  | 20 min – 60 min | < 0.0001 (****) |  |  |
| Malate | 0 min – 20 min | < 0.0001 (****) | P value < 0.0001; R squared 0.9496 |  |
|  | 0 min – 60 min | < 0.0001 (****) |  |  |
|  | 20 min – 60 min | 0.0002 (***) |  |  |

**Supplementary Table 9:**
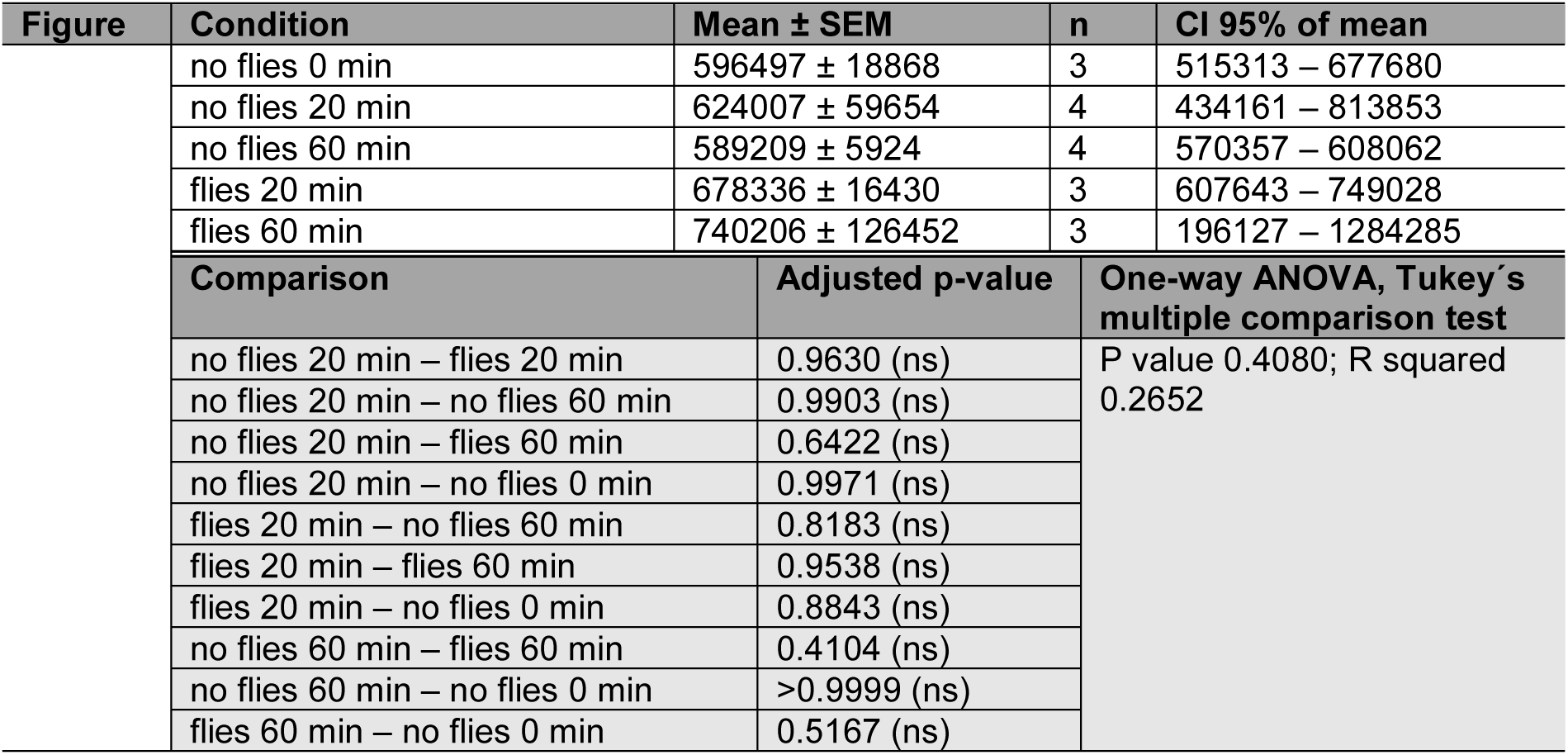
To Supplementary Figure 8

| Figure | Condition | Mean ± SEM | n | CI 95% of mean |
| --- | --- | --- | --- | --- |
| | no flies 0 min | 596497 ± 18868 | 3 | 515313 – 677680 |
| | no flies 20 min | 624007 ± 59654 | 4 | 434161 – 813853 |
| | no flies 60 min | 589209 ± 5924 | 4 | 570357 – 608062 |
| | flies 20 min | 678336 ± 16430 | 3 | 607643 – 749028 |
| | flies 60 min | 740206 ± 126452 | 3 | 196127 – 1284285 |
| Comparison |  | Adjusted p-value | One-way ANOVA, Tukey's multiple comparison test |  |
| no flies 20 min – flies 20 min |  | 0.9630 (ns) | P value 0.4080; R squared 0.2652 |  |
| no flies 20 min – no flies 60 min |  | 0.9903 (ns) |  |  |
| no flies 20 min – flies 60 min |  | 0.6422 (ns) |  |  |
| no flies 20 min – no flies 0 min |  | 0.9971 (ns) |  |  |
| flies 20 min – no flies 60 min |  | 0.8183 (ns) |  |  |
| flies 20 min – flies 60 min |  | 0.9538 (ns) |  |  |
| flies 20 min – no flies 0 min |  | 0.8843 (ns) |  |  |
| no flies 60 min – flies 60 min |  | 0.4104 (ns) |  |  |
| no flies 60 min – no flies 0 min |  | >0.9999 (ns) |  |  |
| flies 60 min – no flies 0 min |  | 0.5167 (ns) |  |  |

## Author Contributions

S.P. designed the research. A.M-E performed the experiments and statistical analyses. B.G. assisted with plates preparation. F.D. and P.G. performed the metabolite measurements and analysis. C.M. and C.L. performed the proteome and acetylome measurements and analysis.

J.T.H. supervised the Seahorse based experiments and assisted with the oxygen measurements. A.M-E. and S.P. wrote the manuscript with input from the other authors. All authors have read and agreed to the published version of the manuscript.

## Conflict of interest

Shahaf Peleg is a co-founder of Luminova Biotech.

## Funding

SP lab is supported by the FBN, DFG grant (458246576), and by two Longevity Impetus grants from Norn Group and by a Longevity Impetus Grant from Norn Group, Hevolution Foundation and Rosenkranz Foundation. BG has been supported by a Longevity Impetus grant from Norn Group. JTH is supported by the DFG CRC1052 (209933838; C7), the free-state of Saxony and Helmholtz Munich.

## Acknowledgments

We would like to thank our technicians Verena Hofer-Pretz and Paula Becker for performing many of the experiments for this study, as well as for managing the laboratory conditions that enabled this work. Further, the authors thank Franziska Hackbarth and Nina Lomp for technical assistance at the BayBioMS and acknowledge Miriam Abele for support in mass spectrometeric data acquisiton.

## Data availability

The mass spectrometric raw files as well as the MaxQuant output files have been deposited to the ProteomeXchange Consortium via the PRIDE partner repository and can be accessed using the identifier PXD050591 (https://proteomecentral.proteomexchange.org/cgi/GetDataset?ID=PXD050591, reviewer account username, password: SMEZOwcq).

